# Structure, dynamics, and inhibition of *Staphylococcus aureus* m^1^A22-tRNA methyltransferase

**DOI:** 10.1101/2021.12.24.474102

**Authors:** Pamela Sweeney, Ashleigh Crowe, Abhishek Kumar, Dinesh Raju, Naveen B. Krishna, Emmajay Sutherland, Caitlin J. Leo, Gemma Fisher, Roopa Lalitha, Likith Muthuraj, Gladstone Sigamani, Verena Oehler, Silvia Synowsky, Sally L. Shirran, Tracey M. Gloster, Clarissa M. Czekster, Pravin Kumar, Rafael G. da Silva

**Affiliations:** School of Biology, Biomedical Sciences Research Complex, University of St Andrews, St Andrews KY16 9ST, UK; Kcat Enzymatic Private Limited, Bangalore, India

**Keywords:** RNA methyltransferase, Transfer RNA (tRNA), *S*-adenosylmethionine (SAM), *Staphylococcus aureus*, X-ray crystallography, TrmK

## Abstract

The enzyme m^1^A22-tRNA methyltransferase (TrmK) catalyses the transfer of a methyl group from SAM to the N1 of adenine 22 in tRNAs. TrmK is essential for *Staphylococcus aureus* survival during infection, but has no homologue in mammals, making it a promising target for antibiotic development. Here we describe the structural and functional characterisation of *S. aureus* TrmK. Crystal structures are reported for *S. aureus* TrmK apoenzyme and in complexes with SAM and SAH. Isothermal titration calorimetry showed that SAM binds to the enzyme with favourable but modest enthalpic and entropic contributions, whereas SAH binding leads to an entropic penalty compensated by a large favourable enthalpic contribution. Molecular dynamics simulations point to specific motions of the C-terminal domain being altered by SAM binding, which might have implications for tRNA recruitment. Activity assays for *S. aureus* TrmK-catalysed methylation of WT and position 22 mutants of tRNA^Leu^ demonstrate that the enzyme requires an adenine at position 22 of the tRNA. Intriguingly, a small RNA hairpin of 18 nucleotides is methylated by TrmK depending on the position of the adenine. *In-silico* screening of compounds suggested plumbagin as a potential inhibitor of TrmK, which was confirmed by activity measurements. Furthermore, LC-MS indicated the protein was covalently modified by one equivalent of the inhibitor, and proteolytic digestion coupled with LC-MS identified Cys92, in the vicinity of the SAM-binding site, as the sole residue modified. These results these results identify a cryptic binding pocket of *S. aureus* TrmK and lay the foundation for future structure-based drug discovery.

## Introduction

Enzyme-catalysed post-transcriptional modifications of tRNA occur in all domains of life, contributing to tRNA stability and folding, proper aminoacylation, and translational fidelity. Among these modifications, methylation is the most abundant and diverse, occurring on both the ribose and nucleobase moieties of tRNA nucleosides at several positions (1–3). All methylation of tRNA adenine N1 (m^1^A) is found in the core region of the tRNA: m^1^A9, m^1^A14, and m^1^A22 in the D-arm, and m^1^A58 in the T-arm (4). The m^1^A22 modification is present only in bacterial tRNA, and was first detected in tRNA^Tyr^ from *Bacillus subtilis* (5). While the biological role of this modification is still elusive, the enzyme responsible for it was identified in 2008, namely m^1^A22-tRNA methyltransferase (TrmK) (6).

TrmK catalyses the transfer of a methyl group from SAM to tRNA, producing m^1^A22-tRNA and SAH (6) (Figure 1). The enzyme belongs to the Class-I methyltransferase family comprised of an N-terminal domain with a Rossmann-like fold for SAM binding and a C-terminal domain with a novel 4-helix fold (7). The latter is analogous to the tRNA-interacting C-terminal domain of the archaeal m^1^G37-tRNA methyltransferase Trm5 (8), and is proposed to play a similar role in TrmK (9).

**Figure 1.**
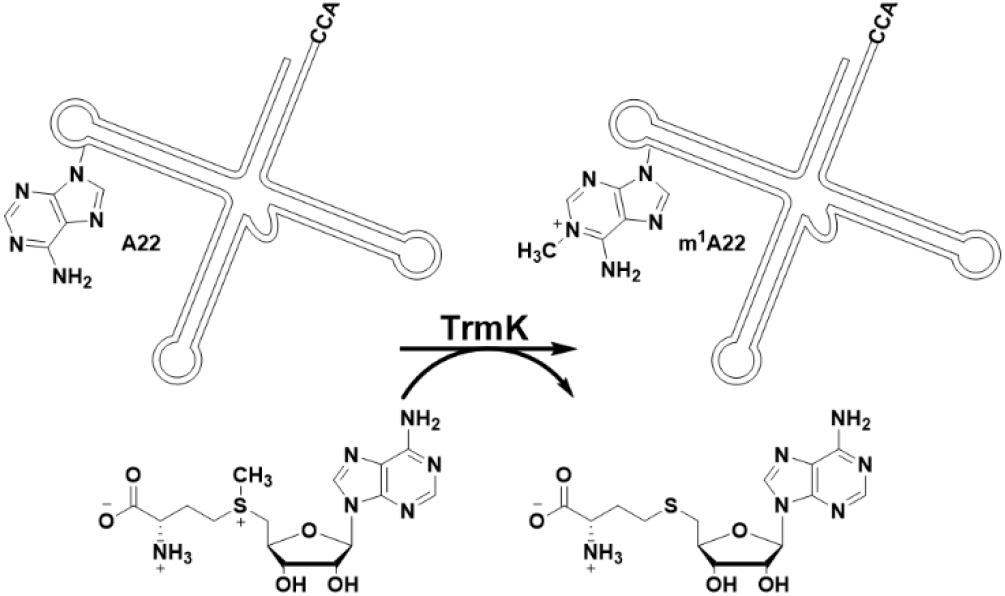
TrmK-catalysed SAM-dependent methylation of N1 of adenine at position 22 of tRNA. Only a schematic rendering of the tRNA is shown.

While TrmK is not required for growth in *B. subtilis* (6), it is essential for survival and growth of the pathogenic Gram-positive bacteria *Staphylococcus aureus* (10–12) and *Streptococcus pneumoniae* (13). Importantly, disruption of the TrmK-encoding gene prevents *S. aureus* survival not only in rich media, but during infection of the bloodstream, vitreous fluid, and cutaneous abscesses in mice (10). Moreover, the TrmK-encoding gene is expressed almost constitutively in cell models of infection (14), during acute and chronic osteomyelitis (15), and in the highly infectious and multidrug-resistant USA300 strain of *S. aureus* during human cutaneous abscess and mouse kidney infection (16). Crucially, TrmK has no homologue in mammals (6), making TrmK a promising target for the development of novel antibiotics against *S. aureus.* This bacterium is a leading cause of healthcare- and community-associated infections worldwide, with many patients infected with methicillin-resistant *S. aureus* (MRSA). MRSA encompasses strains resistant to most antibiotics in clinical use (17). In Europe, MRSA infections are responsible for an average of 7,000 deaths a year, the second highest among all drug-resistant bacterial infections (18), and in the USA, more deaths occur due to MRSA infection than to HIV/AIDS and tuberculosis combined (19). The World Health Organisation ranked MRSA as a high priority in its list of bacteria against which novel antibiotics are urgently needed (20).

In the present work, we have cloned and expressed the gene encoding *S. aureus* TrmK (*Sa*TrmK), purified the recombinant protein, and used X-ray crystallography to obtain high-resolution structures of the enzyme. We employed isothermal titration calorimetry (ITC) and differential scanning fluorimetry (DSF) to characterise cofactor binding, and performed molecular dynamics (MD) simulations to assess differences in flexibility upon ligand binding. We used a luminescence-based SAH-detection assay and LC-MS to uncover the determinants of *Sa*TrmK activity towards tRNA and small RNA hairpins. Finally, we carried out *in-silico* screening of compounds and identified a site on *Sa*TrmK amenable to covalent inhibition.

## Results

### *Sa*TrmK purification and biophysical characterisation

*Sa*TrmK was successfully purified to homogeneity with no other bands visible on a Coomassie Blue-stained gel following SDS-PAGE (Figure S1). A typical yield was 100 mg of *Sa*TrmK per litre of culture. The molecular mass of the protein was confirmed by electrospray-ionisation (ESI)/TOF MS to be 25,564.5 (Figure S2), in agreement with the predicted molecular mass of 25,565.3. DSF assays yielded, upon data fitting to equation 1, a melting temperature (*T*_m_) of 40.3 ± 0.1 °C, which increased modestly to 41.4 ± 0.1 °C and more significantly to 45.9 ± 0.1 °C in the presence of SAH and SAM, respectively (Figure S3). These values suggest SAH binding has little effect on enzyme thermostability. SAM, however, can thermally stabilise the protein upon binding. The *Sa*TrmK elution profile from an analytical gel-filtration column revealed a major peak at 16 mL (MW of ~23 kDa) and a minor peak at 14.4 mL (MW of ~50 kDa) (Figure S4). This profile indicates most of the protein is monomeric, while a minor fraction is dimeric.

### Crystal structures of *Sa*TrmK apoenzyme, *Sa*TrmK:SAM and *Sa*TrmK:SAH

To shed light on the structural basis for cofactor binding and help inform future structure-based inhibitor design, crystals of *Sa*TrmK apoenzyme and the binary complexes *Sa*TrmK:SAM and *Sa*TrmK: SAH were obtained and data were collected to a resolution of 1.1, 1.4, and 1.5 Å, respectively (Table S1). The 2m*F*_0_ - D*F*_c_ map showed electron density for SAM and SAH in the respective structures (Figure 2). *Sa*TrmK is typical of Class-I methyltransferases, with an N-terminal domain where the cofactor binds and methyl transfer takes place, and a C-terminal domain likely involved in tRNA recognition (8,9). Superposition of the three structures (Figure S5) revealed negligible effect of cofactor binding on Cα root-mean-square deviations (RMSD), whose values were 0.15, 0.18, and 0.08 Å between *Sa*TrmK and *Sa*TrmK:SAM, *Sa*TrmK and *Sa*TrmK:SAH, and *Sa*TrmK:SAM and *Sa*TrmK:SAH, respectively. The *Sa*TrmK apoenzyme structure superposes with the apoenzyme structures of *B. subtilis* TrmK (PDB entry 6Q56) (9) and *S. pneumoniae* (PDB entry 3KR9) (7) with Cα RMSDs of 0.68 and 0.98 Å, respectively, with the most ostensible differences located in the coiled-coil motif of the C-terminal domain (Figure S6).

**Figure 2.**
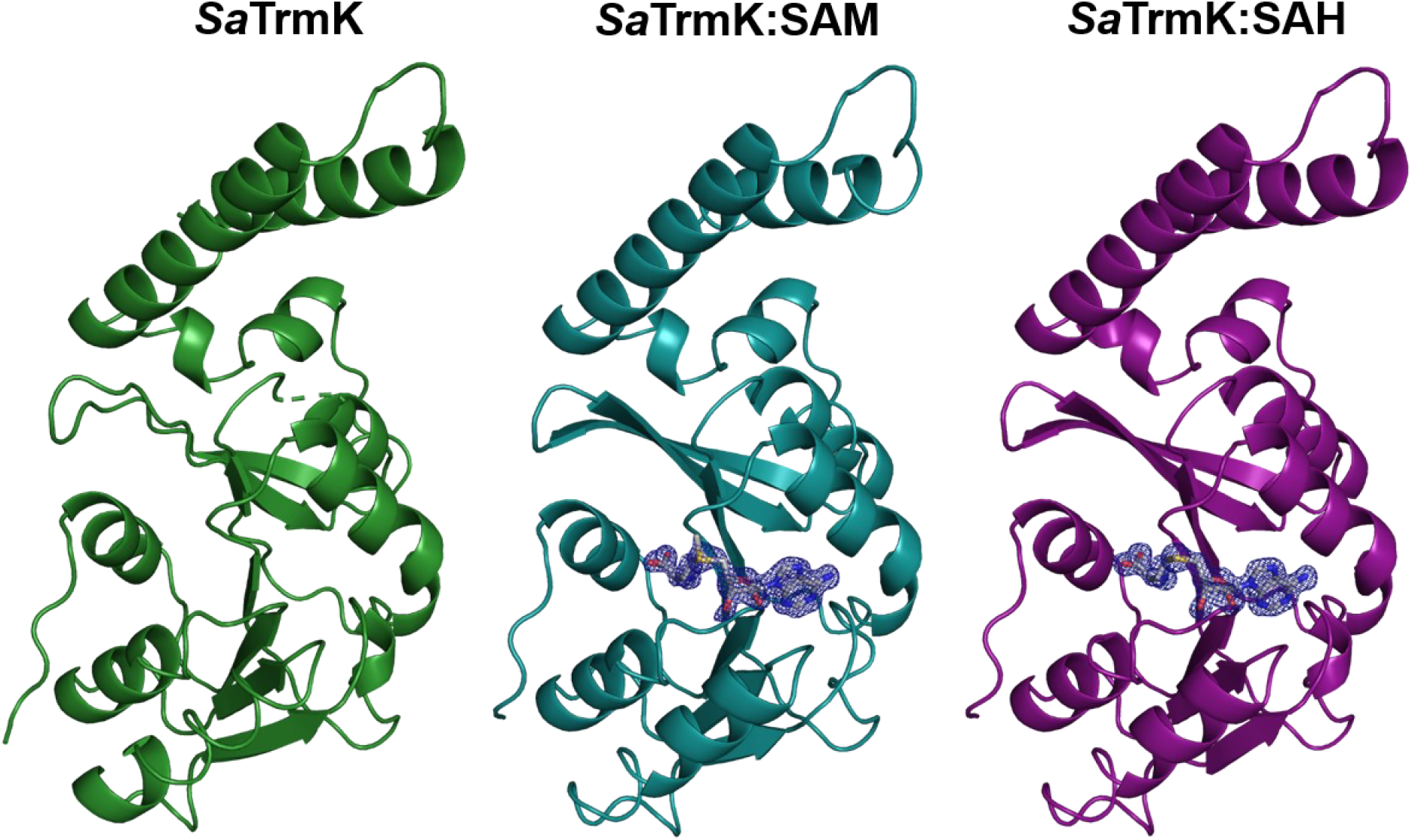
Ribbon diagram of the crystal structures of *Sa*TrmK apoenzyme and the binary complexes with SAM and SAH. Electron density for SAM and SAH (2m*F*_o_ – D*F*_c_ maps at 1σ) is shown in blue. The electron density for SAM is partially disordered at the methyl which is transferred. The cofactors are shown in stick models, with oxygen in red, nitrogen in blue, carbon in grey, and sulphur in yellow.

The electron density data for the *Sa*TrmK apoenzyme showed clearly defined density near the active site that could not be fitted with any of the small molecules involved in purification or crystallisation of the enzyme, suggesting it was carried through with *Sa*TrmK from protein expression. Intriguingly, citrate could be fitted perfectly into the electron density, making polar contacts with the side chains of His27, Tyr29, and Asn59 (Figure S7A). It is possible the negatively charged groups of citrate is mimicking the phosphate group(s) of tRNA. DSF analysis indicated that citrate leads to an overall modest decrease in *Sa*TrmK *T_m_* (Figure S7B) that only becomes noticeable at high citrate concentrations, which is incompatible with the presumed tight binding required for the interaction to continue from protein expression to crystallisation. Therefore, the significance of this observation, if any, is still elusive.

### The cofactor binding site

The binding site for SAM is restricted to the N-terminal domain, and several polar and nonpolar interactions are made with residues via both main- and side-chain moieties to position the cofactor in the active site (Figure 3A). SAM’s COO^−^ group forms a salt bridge with the ω-NH_2_ and ω′-NH_2_ groups of Arg7, while its NH_3_+ group donates a hydrogen bond (H-bond) to the main-chain CO-groups of Cys92 and Gly24, and to a water molecule (W1) which in turn H-bonds to Asp22. The 2’-OH and 3’-OH form H-bonds with another water molecule (W2) that itself H-bonds to Glu47 side-chain COO^−^ and Val48 main-chain –NH. Adenine N1 and N6 atoms form H-bonds with Gly76 main-chain –NH and Asp75 side-chain COO^−^, respectively. Furthermore, the adenine plane is flanked on both sides by hydrophobic interactions with Met94, Leu98, Ile102 and Val48.

**Figure 3.**
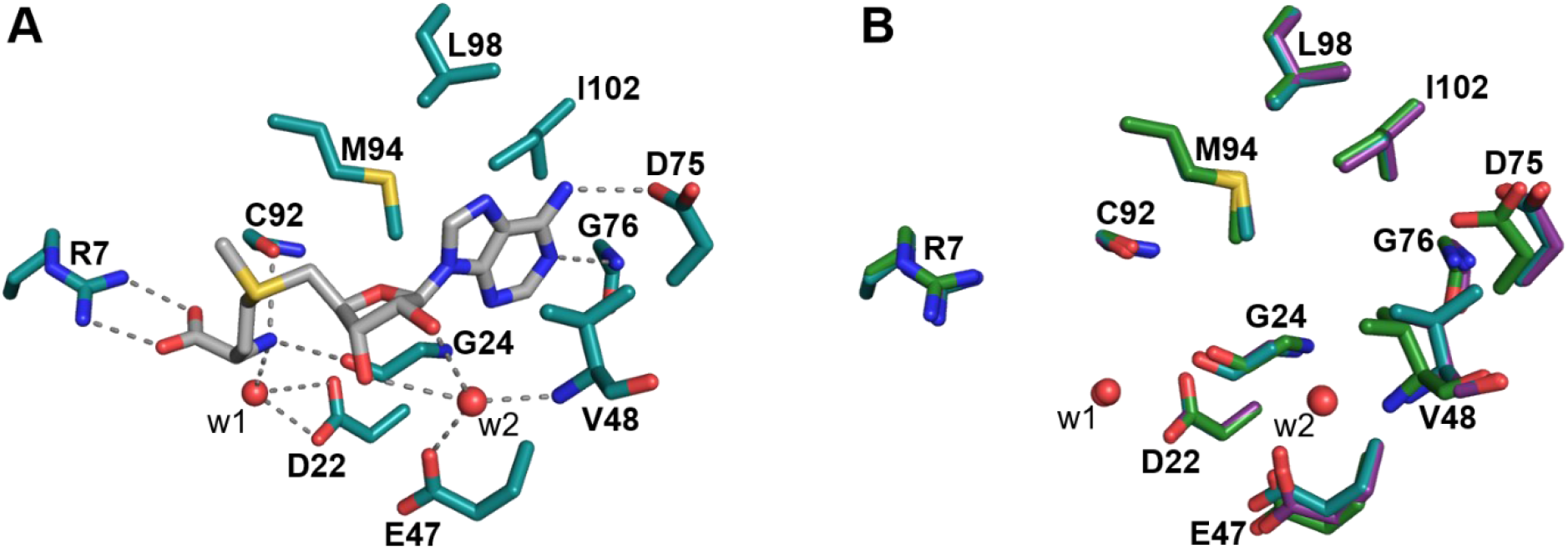
*Sa*TrmK cofactor binding site. (A) Key polar and nonpolar interactions between *Sa*TrmK residues and SAM. The dashed lines depict polar contacts. (B) Overlay of the cofactor-binding site of *Sa*TrmK apoenzyme and the binary complexes with SAM and SAH. In both panels, oxygen is depicted in red, nitrogen in blue, sulphur in yellow, and carbon in either grey (SAM), teal (residues in *Sa*TrmK:SAM), purple (residues in *Sa*TrmK:SAH), or green (residues in *Sa*TrmK apoenzyme). Water molecules are shown as red spheres.

A superposition of the active sites of *Sa*TrmK, *Sa*TrmK:SAM, and *Sa*TrmK:SAH (Figure 3B) shows there is local motion of some residues in order to accommodate the cofactor. In the region of the active site which binds the adenine moiety of SAM, the side chain of Glu47 moves slightly from its position in the apoenzyme but retains the same orientation. Interestingly, water molecule W2 is only observed in the structures when SAM and SAH are bound. Val48 moves ~1 Å and rotates approximately 120° enabling both methyl groups to face away from SAM, and Asp75 also moves ~1 Å, and rotates slightly. Their position in the apoenzyme would have caused a steric clash with SAM. Asp26, which is in the vicinity of the SAM binding site but does not interact with the cofactor, moves significantly upon cofactor binding from its position in the apoenzyme (Figure S8).

### *Sa*TrmK binds SAM and SAH with distinct thermodynamics

In order to confirm binary complex formation in solution as observed *in crystallo* and gain insight into cofactor binding equilibrium thermodynamics, SAM and SAH binding to *Sa*TrmK were determined by ITC (Figure 4). The binding isotherms were best fitted to a single-site binding model with 1:1 (protein:ligand) stoichiometry. Binding of SAM to *Sa*TrmK yielded a *K*_D_ of 39 ± 4 μM, while binding of SAH resulted in a *K*_D_ of 0.94 ± 0.05 μM. The Gibbs free energies (ΔG) were −5.92 ± 0.07 and −8.09 ± 0.03 kcal/mol for SAM and SAH binding to *Sa*TrmK, respectively, with SAH binding favoured by 2.17 ± 0.08 kcal/mol. Interestingly, while binding of both SAM and SAH are exothermic, they are driven by distinct thermodynamic functions. The *Sa*TrmK:SAM complex is stabilised by modest but favourable enthalpic and entropic contributions, with ΔH of −3.46 ± 0.08 kcal/mol and TΔS of 2.5 ± 0.1 kcal/mol. On the other hand, the *Sa*TrmK: SAH complex formation is enthalpically driven, with ΔH of −12.8 ± 0.2 kcal/mol which pays for an entropic penalty on binary complex formation, given a TΔS of −4.7 ± 0.3 kcal/mol.

**Figure 4.**
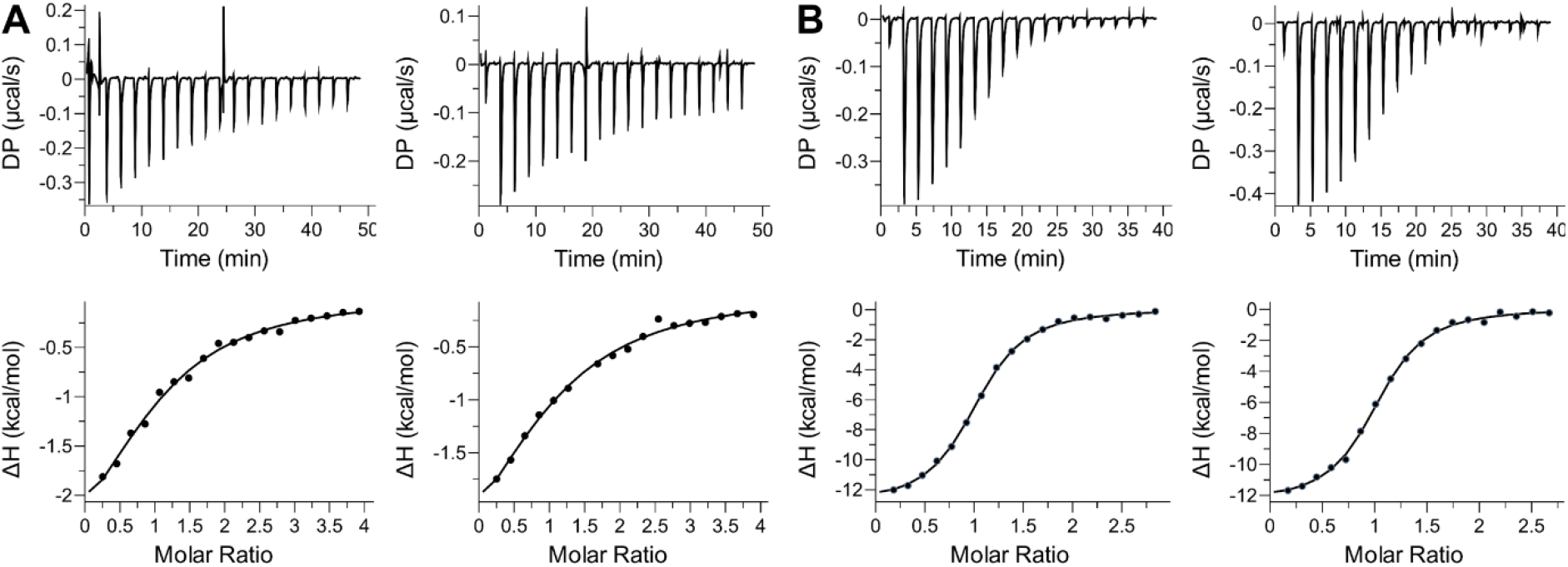
Characterisation of cofactor binding to *Sa*TrmK by ITC. (A) Two independent titrations of *Sa*TrmK with SAM. The 7^th^ injection on the graph on the top-right panel was an outlier and was excluded from the fit in the bottom-right panel. (B) Two independent titrations of *Sa*TrmK with SAH. In all cases, the data were best fitted to a single-site binding model with 1:1 (protein:ligand) stoichiometry.

### SAM binding alters specific motions of *Sa*TrmK

To gain insight into the dynamics of *Sa*TrmK, MD simulations were carried out over 500 ns using the crystal structures as starting points (Figure 5). The distance of the salt bridge between Arg7 and either SAM or SAH confirmed the ligands did not dissociate from the protein over the course of the simulations but showed that the salt bridge with SAM oscillated more in comparison with SAH (Figure 5A). Time-dependent Cα RMSDs indicated only minor oscillations for the three structures (Figure 5B), and the overall root-mean-square fluctuations (RMSFs) were remarkably similar, with the most flexible region comprising the C-terminal domain (Figure 5C). Moreover, correlated motions between protein domains are similar among the three structures (Figure S9).

**Figure 5.**
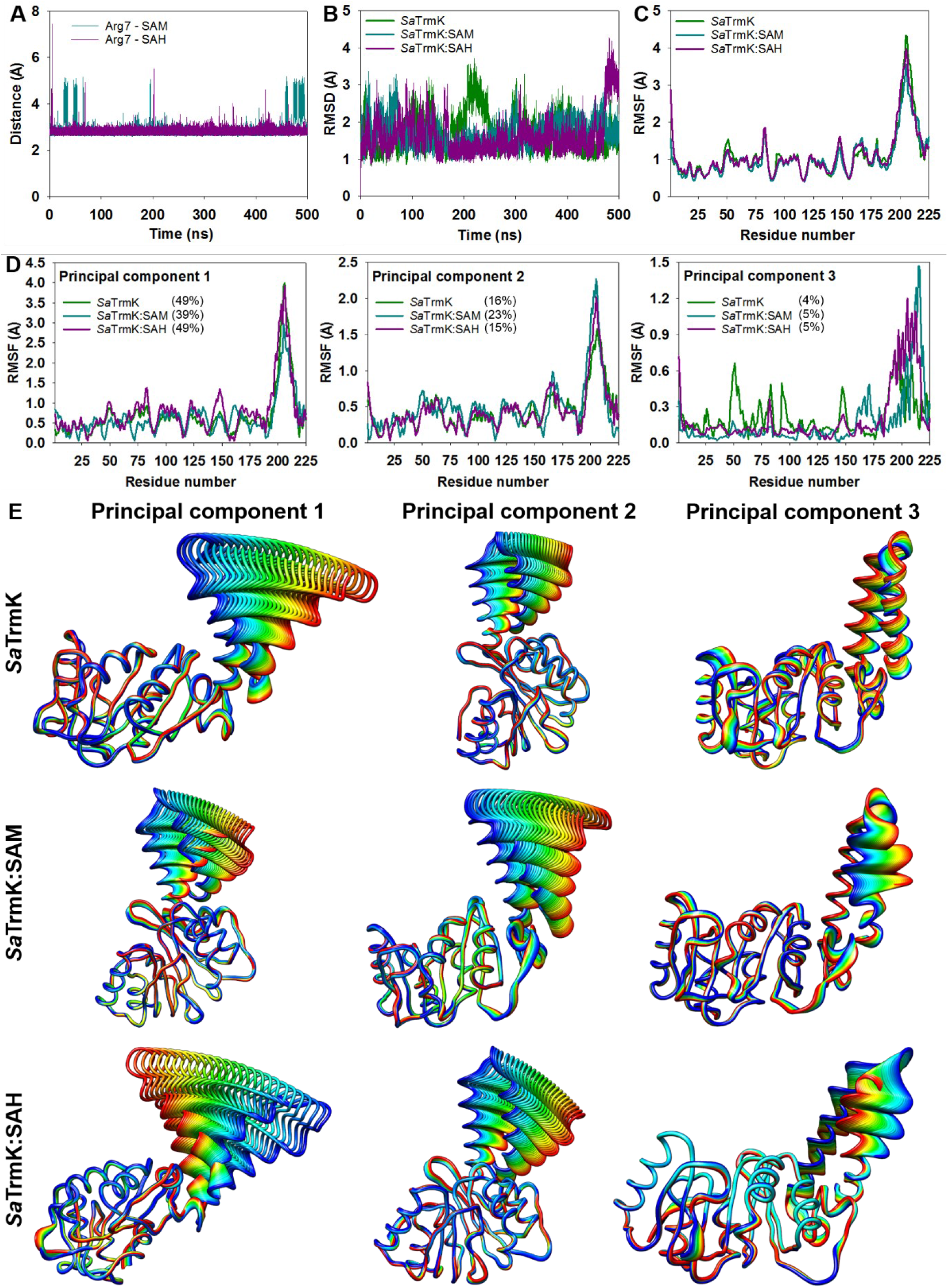
MD simulations of *Sa*TrmK. (A) Mean length of the salt bridge between Arg7 and either SAM or SAH. (B) Time-dependence of the RMSDs over *Sa*TrmK, *Sa*TrmK:SAM, and *Sa*TrmK:SAH Ca. (C) Mean RMSF over Ca. (D) Ca RMSFs of the top-three dominant motions of *Sa*TrmK, *Sa*TrmK: SAM, and *Sa*TrmK: SAH as revealed by principal component analysis. Percentages in brackets represent the relative eigenvalue contributed by each eigenvector in each structure. (E) Ribbon-diagram representation of each dominant motion. The width or delocalisation of the ribbon regions corresponds to the motion amplitude.

In order to uncover dominant motions of *Sa*TrmK, principal component analysis was carried out, and the first three eigenvectors were isolated (Figure 5D). This analysis revealed that the first two eigenvectors are dominated by specific motions of the C-terminal domain, but SAM binding is accompanied by a swap in the specific trajectories contributing to the highest RMSFs, as compared with *Sa*TrmK apoenzyme and *Sa*TrmK:SAH binary complex (Figure 5E). Furthermore, the third eigenvector suggests that SAM and SAH binding reduce the amplitude of specific motions in the N-terminal domain, most notably those around residues implicated in cofactor binding, while increasing specific motions in the C-terminal domain, as compared with the apoenzyme (Figure 5D and E).

Molecular electrostatic potential surfaces were calculated for the three structures throughout the simulations, and snapshots were taken every 125 ns (Figure S10). While the cofactor binding pocket is consistently dominated by negative charges in all structures, positive charges outside this pocket undergo some redistribution throughout the trajectories, influenced by the presence or absence of SAM or SAH.

### *Sa*TrmK-catalysed methylation of tRNA^Leu^

The MTase-Glo™ Methyltransferase Assay, which detects luminescence generated in a luciferase-coupled reaction driven by ATP, which in turn is synthesized enzymatically from the SAH produced in the methyltransferase reaction (21), was used to quantify the methylation of *S. aureus* tRNA^Leu^ as catalysed by *Sa*TrmK. This is a robust and sensitive assay used to characterise RNA methyltransferase activity (22) and to screen tRNA methyltransferase inhibitors (23), as it has been shown to result in lower false-positive rates than other methods (24). In the absence of tRNA, *Sa*TrmK contributes no background luminescence, as expected since the enzyme is not purified bound to SAH. This is evidenced by the crystal structure of the apoenzyme and no SAH detected by LC-MS after denaturation of *Sa*TrmK which would have released any bound SAH (Figure S11). Background luminescence possibly originating from contaminating ATP in the RNA samples was drastically diminished by dialysis of both tRNA and small RNA hairpins prior to use, and any residual background luminescence was subtracted from the reactions via controls lacking RNA. An SAH standard curve (Figure S12) was used for quantification of the methylation reaction.

Activity of *Sa*TrmK was detected when tRNA^Leu^, which contains an adenine at position 22, was used as substrate, but no SAH formation could be detected when either A22C-tRNA^Leu^, A22U-tRNA^Leu^, or A22G-tRNA^Leu^ was tested as substrate (Figure 6A), suggesting methylation is conditional on an adenine at position 22. Furthermore, SAH production was linearly dependent on *Sa*TrmK concentration with tRNA^Leu^ as substrate (Figure 6A, inset). To validate the assay further for detection of *Sa*TrmK enzymatic activity, SAH production was measured in the presence and absence of sinefungin, a SAM analogue that is a pan-methyltransferase inhibitor (25). Sinefungin led to ~98% inhibition of SAH production (Figure 6B). In the absence of *Sa*TrmK, sinefungin had no effect on luminescence generated by the addition of SAH to the assay (Figure S13), pointing to sinefungin preventing SAH production by inhibiting *Sa*TrmK. When the tRNA^Leu^ was hydrolysed to nucleoside 5’-monophosphates by nuclease P1 following the methylation reaction, and subjected to LC-MS analysis, ions with the same mass/charge (m/z) and retention time as those for the N^1^-methyl-AMP standard were detected (Figure 6C). However, no such signal was detected when either *Sa*TrmK was omitted from the methylation reaction or A22C-tRNA was used instead of tRNA^Leu^ (Figure 6C). This is in strict accordance with TrmK producing m^1^A22-tRNA (6). Moreover, by directly detecting the methylated adenine, the LC-MS-based assay provided orthogonal validation of the MTase-Glo™ Methyltransferase Assay for detection of *Sa*TrmK activity.

**Figure 6.**
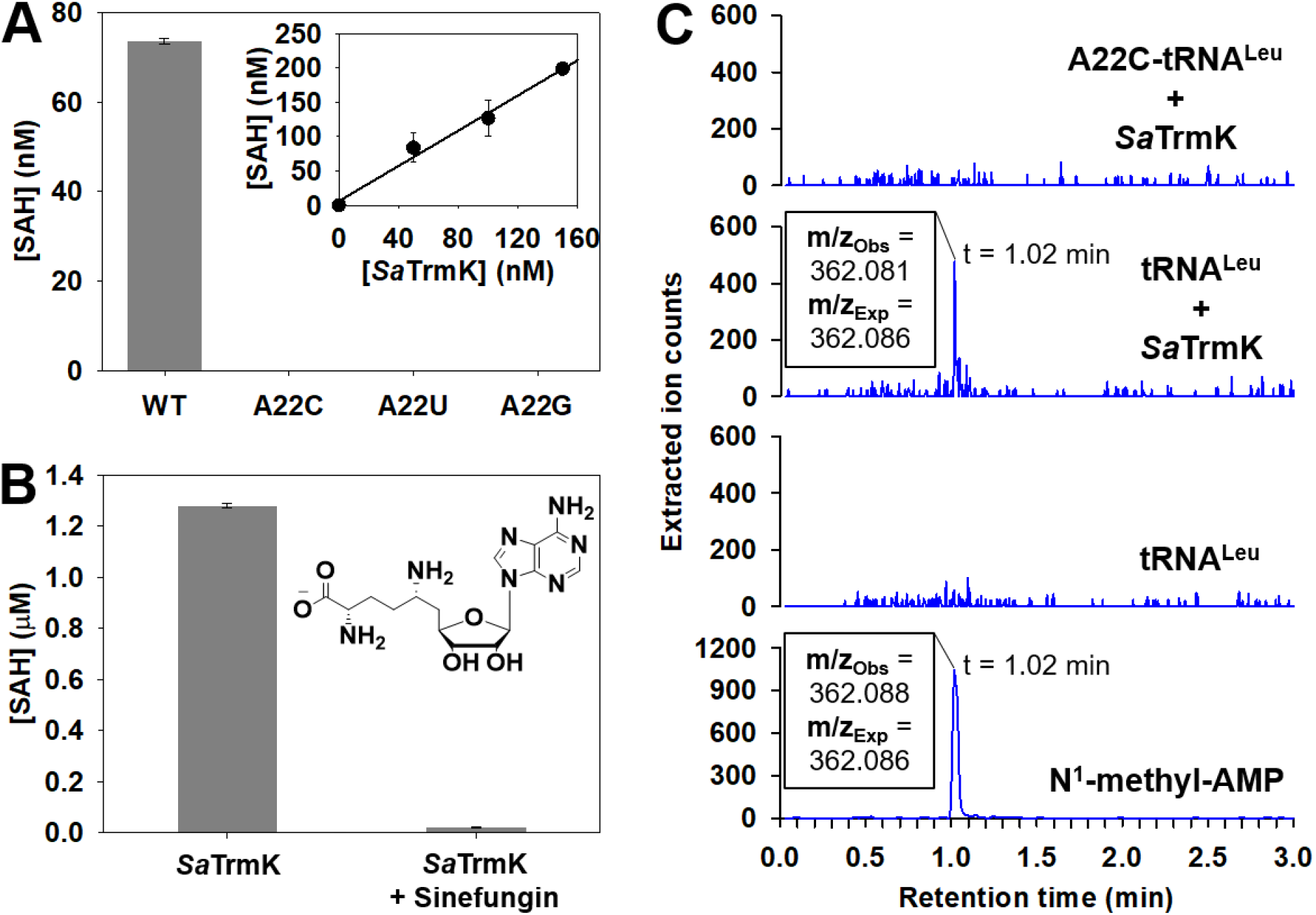
*Sa*TrmK enzymatic activity with *S. aureus* tRNA^Leu^. (A) Methylation of WT and mutant tRNA^Leu^ by *Sa*TrmK. The inset shows the dependence of SAH formed during the methylation of tRNA^Leu^ on *Sa*TrmK concentration. (B) Inhibition of *Sa*TrmK by sinefungin. The inset depicts the chemical structure of sinefungin. (C) LC-MS analysis of *Sa*TrmK-catalysed methylation of tRNA^Leu^. The panel labelled N^1^-methyl-AMP refers to the commercial compound used as a standard.

### *Sa*TrmK-catalysed methylation of short RNA hairpins

*B. subtilis* TrmK has been reported to require a full-length tRNA for activity, since no methyl transfer was detected when smaller RNAs were tested as substrate (9). To confirm the same was true for *Sa*TrmK, an RNA^18mer^, predicted to form a hairpin similar to the D-arm of tRNA but with only 4 nucleotides in the loop and containing adenine at three positions (Figure 7A), was designed and tested as a substrate of the enzyme. Surprisingly, *Sa*TrmK accepted this RNA^18mer^ as a substrate (Figure 7A). Catalytic activity increased to a level approaching that detected with tRNA^Leu^ (Figure 6A) when A10G-RNA^18mer^ replaced the original RNA^18mer^ (referred to as WT-RNA^18mer^), but it was undetectable with either A11G-RNA^18mer^ or U5C/A14G-RNA^18mer^ as the substrate (Figure 7A). This suggests that A10G-RNA^18mer^ can be efficiently methylated by *Sa*TrmK, pointing to a novel property of this enzyme.

**Figure 7.**
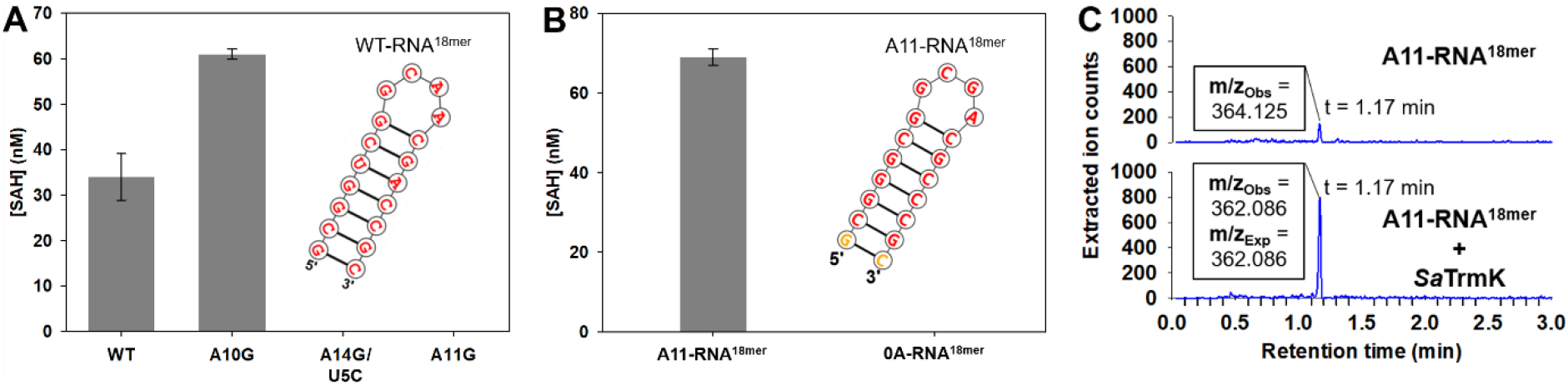
*Sa*TrmK enzymatic activity with RNA^18mer^. (A) Methylation of WT and mutant RNA^18mer^ by *Sa*TrmK. The inset shows the nucleotide sequence and predicted secondary structure of the WT-RNA^18mer^. (B) Methylation of A11-RNA^18mer^ and 0A-RNA^18mer^ by *Sa*TrmK. The inset shows the nucleotide sequence and predicted secondary structure of the A11-RNA^18mer^. (C) LC-MS analysis of *Sa*TrmK-catalysed methylation of A11-RNA^18mer^.

The lack of methylation activity towards U5C/A14G-RNA^18mer^ was puzzling since A14 should be in a Watson-Crick pair with U5 in the WT-RNA^18mer^, making its N1 unavailable as a nucleophile for the methyl-transfer reaction, and A11 would be the only expected methylation site in the A10G-RNA^18mer^. Hence it is possible the C5 and G14 substitution caused a decrease in affinity for *Sa*TrmK and/or reduction of A11 methylation efficiency. To probe the limits of RNA^18mer^ methylation further, a second-generation RNA^18mer^ was synthesised which contained G5, C14, and only one adenine, at position 11 (A11-RNA^18mer^). At higher concentrations of RNA and *Sa*TrmK than previously used, significant methylation of A11-RNA^18mer^ was detected (Figure 7B). To confirm SAH formation was indeed a result of *Sa*TrmK-catalysed methylation of A11 in A11-RNA^18mer^ and not an unforeseen artefact of the higher substrate and enzyme concentrations, a no-adenine (0A-RNA^18mer^) counterpart to A11-RNA^18mer^ harbouring an A11G substitution was tested as a substrate for *Sa*TrmK under the same conditions as A11-RNA^18mer^, and no activity was detected (Figure 7B). Again, orthogonal evidence for methylation of A11-RNA^18mer^ was obtained when A11-RNA^18mer^ was hydrolysed to nucleoside 5’-monophosphates by nuclease P1 following the methylation reaction, and subsequently subjected to LC-MS analysis. Ions were detected with the exact m/z (362.086) expected for N^1^-methyl-AMP. However, no such signal was detected in a control sample where *Sa*TrmK was omitted from the methylation reaction. A small peak rising slightly above the noise level was detected in the control with the same retention time as the presumed N^1^-methyl-AMP peak from the reaction sample, but with an m/z = 364.125 that does not correspond N^1^-methyl-AMP (Figure 7C).

### Covalent inhibition of *Sa*TrmK by plumbagin

In order to identify potential inhibitors of *Sa*TrmK, *in-silico* screening of compounds based on molecular docking was carried out against the *Sa*TrmK:SAM binary complex structure. This complex was chosen to increase the stringency of the screening against compounds that would compete with SAM for the cofactor-binding site, in an attempt to discover potentially novel druggable pockets in *Sa*TrmK while increasing selectivity against the myriad other SAM-dependent methyltransferases in humans. The top hits from the screening were tested as inhibitors of *Sa*TrmK, however, all but one, plumbagin (5-hydroxy-2-methyl-1,4-naphtoquinone), significantly interfered with the activity assay itself and were discarded. Plumbagin docked to a pocket adjacent to the SAM-binding site, making nonpolar contacts with Ile20 and Leu30 and polar interactions with Asp22 and Asp26 (Figure 8A). Plumbagin led to ~93% inhibition of *Sa*TrmK-catalysed methylation of tRNA^Leu^ (Figure 8B). In the absence of *Sa*TrmK, plumbagin had a negligible effect (~6% decrease) on luminescence generated by the addition of SAH to the assay (Figure S14).

**Figure 8.**
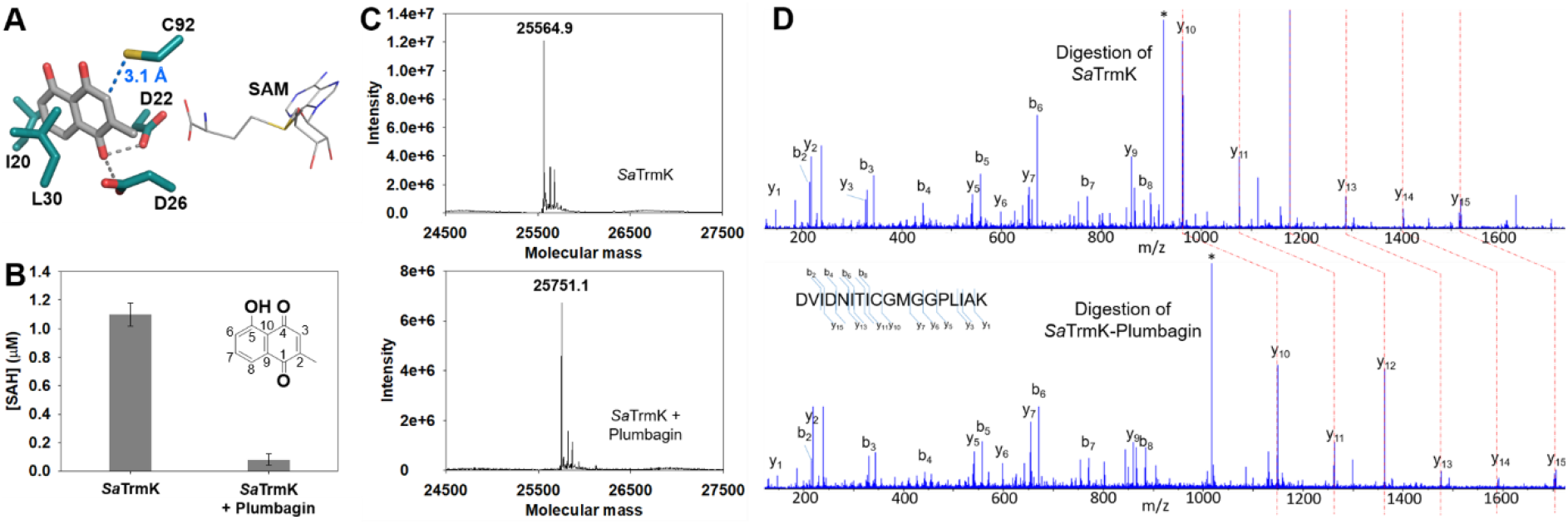
Inhibition of *Sa*TrmK by plumbagin. (A) of the molecular docking-predicted binding of plumbagin to an *Sa*TrmK site adjacent to the SAM-binding site. Plumbagin and protein are shown in stick model, while SAM is shown in wireframe. Oxygen is depicted in red, nitrogen in blue, sulphur in yellow, and carbon in either grey (SAM and plumbagin) or teal (*Sa*TrmK). (B) Inhibition of *Sa*TrmK by plumbagin. The inset depicts the chemical structure of plumbagin with ring carbons numbered. (C) Intact mass of vehicle-treated (top) and plumbagin-treated (bottom) *Sa*TrmK, showing an increase in molecular mass of 186.2 upon incubation with plumbagin. (D) MS/MS fragmentation data for [M+2H]^2+^ = 923.4972 and [M+2H]^2+^ = 1,016.5033 for trypsin/Glu-C-digested vehicle-treated (top) and plumbagin-treated (bottom) *Sa*TrmK. The b- and y-fragmentation ions are indicated, with shift of 186.0122 at y10. The * denotes the unfragmented peptide.

Plumbagin is a Michael acceptor and can undergo conjugate addition with amino and thiolate groups on its C3 position (26,27), and in the docking model the C3 is 3.1 Å away from the thiol group of Cys92 (Figure 8A). Thus, the hypothesis was considered that plumbagin inhibits *Sa*TrmK via covalent attachment to Cys92 to form a Michael adduct. Incubation of *Sa*TrmK with either the vehicle (1% DMSO) or with plumbagin followed by determination of the protein intact mass by ESI-MS yielded molecular mass of 25,564.9 and 25,751.1, respectively, indicating that upon incubation with plumbagin, *Sa*TrmK mass increased by 186.2 (Figure 8C). To identify the specific site of the modification, the samples were digested with trypsin and Glu-C and subjected to LC-MS/MS analysis. Only one peptide, spanning residues Asp84 – Lys101 (NH_2_-DVIDNITICGMGGPLIAK-CO2H) showed a difference in mass between vehicle-treated and plumbagin-treated samples (Figure 8D), detected as [M+2H]^2+^ = 923.4972 and [M+2H]^2+^ = 1,016.5033, respectively (Met94 is oxidised in both samples), which resulted in a mass difference of 186.0122; *de novo* sequencing showed the peptides have the same amino acid backbone, but differ in mass by 186.0122 at Cys92 as indicated by the shift in the y-ion fragmentation series at y10 (Figure 8D). These results confirm that plumbagin inhibits *Sa*TrmK via covalent attachment to Cys92, but the mass difference after the covalent modification is 2 mass units lower than the 188 expected for a Michael adduct (Figure S15A).

## Discussion

*Sa*TrmK is an essential enzyme for *S. aureus* survival and infection, and promising target for the development of novel drugs to treat MRSA infections. The high-resolution structures of *Sa*TrmK presented here along with the functional characterisation of the enzyme provide a starting point for understanding its catalytic mechanism and for designing specific inhibitors. Based on the crystal structures, SAM binding does not lead to significant conformational changes in the enzyme. Nonetheless, the different rotamer adopted by Asp26 upon SAM binding might point to a role for this residue as a general base to abstract a proton from the 6-NH_2_ group of tRNA A22, thus increasing the nucleophilicity of N1. Supporting a role in catalysis, Asp26 is highly conserved among TrmK orthologues; however, in *B. subtilis* TrmK, replacement of the equivalent residue Asp29 for alanine had a detrimental but not completely impairing effect on catalysis (9). Moreover, similar reactions where adenine N1 acts as a nucleophile have been proposed to proceed without general base catalysis (28), likely taking advantage of the transient negative charge on N1 due to a natural resonance structure of adenine (29).

The ability of *Sa*TrmK to bind SAM or SAH in the absence of tRNA, along with the cofactor binding site being located in a cleft in the N-terminal domain with the methyl group of SAM to be transferred pointing outwards, leads to the hypothesis the kinetic mechanism is ordered with SAM binding first to the enzyme and SAH departing last, since binding of the tRNA first would be expected to block access to the cofactor binding pocket. The *K*_D_ of 0.94 μM measured here for the *Sa*TrmK:SAH complex is in agreement with that reported for *B. subtilis* TrmK binding to SAH (1.7 μM) (9), whereas the *K*_D_ of 39 μM for dissociation of the *Sa*TrmK: SAM complex closely matches those reported for SAM binding to *Sulfolobus acidocaldarius* m^1^A9-tRNA methyltransferase (33 μM) (30) and to *S. aureus* rRNA methyltransferase OrfX (52 μM) (31). The ~40-fold lower affinity of *Sa*TrmK for SAM than for SAH is intriguing, given the remarkably similar interactions of both molecules with the active site, and might reflect instead possible differences in desolvation energy between the two molecules.

Molecular dynamics simulations point to the C-terminal domain as the most conformationally flexible region of *Sa*TrmK, in agreement with what was proposed for *B. subtilis* TrmK (9). In the archaeal Class-I methyltransferase Trm5, an analogous, though not homologous, C-terminal domain is involved in tRNA binding away from the methylation site (8), and in *B. subtilis* TrmK, partial deletion of the C-terminal domain generated protein that could bind SAH with unaltered affinity but showed no catalytic activity (9). In *Sa*TrmK, SAM and SAH binding increases the amplitude of specific motions of this domain as compared with the apoenzyme, suggesting that cofactor binding-modulated dynamics of the C-terminal domain might be involved in tRNA binding or m^1^A22-tRNA release.

*Sa*TrmK demonstrates a strict requirement for adenine at position 22 of tRNA for methylation, as exemplified by its inability to methylate A22G-, A22C and A22U-tRNA^Leu^. In *S. aureus*, the m^1^A22 modification has been detected *in vivo* on several tRNAs (32), suggesting the presence of A22, not the identity of the tRNA itself, as the key factor in substrate selectivity by *Sa*TrmK. This is in agreement with data for *B. subtilis* TrmK, where a broad substrate specificity was reported though the presence of G13 was also a requirement along with A22 (9). However, in contrast to the *B. subtilis* orthologue, which requires the full-length tRNA for activity, *Sa*TrmK is able to methylate small RNA hairpins provided they harbour an adenine at certain positions. The relevance *in vivo*, if any, of this finding is yet unknown, but it is in line with reports for other tRNA methyltransferases which have been shown to accept small RNA hairpins as substrates *in vitro*, for instance *E. coli* m^5^U54-tRNA methyltransferase (TrmA) (33) and m^1^G-tRNA methyltransferase (34). At a minimum, this observation may render the RNA^18mers^ used here useful research tools to explore the mechanism of *Sa*TrmK. Co-crystals of methyltransferase:tRNA are notoriously difficult to obtain, given the large size and flexibility of the tRNA, and in the case of TrmA, the small RNA hairpin served as a tool for crystallography, allowing the crystal structure of TrmA:RNA^19mer^ complex to be determined (34).

The very low activity of *Sa*TrmK towards the A11-RNA^18mer^, along with the strict requirement for that sole adenine since no methylation is detectable towards 0A-RNA^18mer^, may provide an ideal substrate for future transition-state analysis from kinetic isotope effects. Kinetic isotope effects are only expressed in reactions where the chemical step is slow in relation to physical steps such as substrate dissociation and conformational changes (35). With enzymes that act on macromolecules, the often observed tight-binding of the full-length substrates usually translates in a very low substrate dissociation rate, but the use of truncated versions of the substrate, for instance with the DNA methyltransferase 1, often leads to high dissociation rates, reduced probability for the chemical step, and significant expression of kinetic isotope effects (36).

Finally, a novel binding site was identified on *Sa*TrmK by molecular docking, and the potential ligand plumbagin was shown to be a covalent inhibitor of the enzyme. The mechanism for covalent adduct formation is unclear. While an initial Michael addition is the most appealing hypothesis given the structure of plumbagin and its position relative to Cys92 as predicted by the docking model, extensive LC-ESI-MS/MS analysis of the *Sa*TrmK-plumbagin adduct yields a mass difference of ~186 instead of 188 as expected for a Michael adduct. One possibility is oxidation of a putative initial Michael adduct (Figure S15B), which would result in the observed mass difference of 186. Michael adducts of plumbagin have been reported to undergo oxidation via an epoxide intermediate involving C2 and C3, for instance in the biosynthesis and organic synthesis of zeylanone (37). Again, the oxidation mechanism the plumbagin binding site on *Sa*TrmK could accommodate is elusive. All that can be stated based on the experimental evidence is that incubation of *Sa*TrmK with plumbagin leads to an inactivated enzyme via covalent modification of Cys92.

Elucidation of the exact mechanism for covalent modification is beyond the scope of this work, since plumbagin is not an attractive *Sa*TrmK inhibitor *per se* in terms of drug development due to its lack of target selectivity, in light of its disrupting the function of several human proteins (38,39). The key relevance of the inhibition of *Sa*TrmK by plumbagin is the discovery of a cryptic pocket on the enzyme where a cysteine lies that is amenable to covalent modification. Cryptic pockets are potential binding sites on a protein that are not apparent from the structure of the apoprotein and usually did not evolve to bind a natural ligand, revealing themselves only when a compound binds to it (40). The plumbagin-binding site of *Sa*TrmK is not where the cofactor binds, and judging by a docking model of tRNA-bound *B. subtilis* TrmK (9), it is not involved in tRNA binding either. This increases the possibility that *Sa*TrmK inhibition targeting this site will not be competitively overcome once the substrates accumulate. Furthermore, the demonstrated feasibility of covalently accessing Cys92 bodes well for increased potency and sustained target engagement of future inhibitors (41). One caveat is that Cys92 is not extensively conserved across TrmK orthologues (9), bringing about the possibility that a single mutation at position 92 could eliminate covalent inhibition without impairing catalytic efficiency. This is scenario is mitigated, however, by the strict conservation of Asp22 and As26, and high conservation of the hydrophobic residues lining the pocket (9), offering opportunities for the design of alternative non-covalent inhibitors of *Sa*TrmK that are specific, potent, and resilient.

## Experimental procedures

### Materials

Nucleoside 5’-triphosphates, MgCl_2_, dithiothreitol (DTT), glycerol, lysozyme, DNAse I, kanamycin, spermidine, plumbagin, sinefungin, SAM and SAH were purchased from Sigma-Aldrich. EDTA-free Cømplete protease inhibitor cocktail tablets were from Roche. Isopropyl *β*-D-1-thiogalactopyranoside (IPTG), HEPES, and NaCl were purchased from Formedium, and N^1^-methyladenoside 5’-phosphate (N^1^-methyl-AMP) was from Jena Bioscience. MTase-Glo™ Methyltransferase Assay Kit containing SAM and SAH were purchased from Promega. All other chemicals were purchased from readily available commercial sources and were used without further purification. Tobacco etch virus protease (TEVP) was produced as previously described (42). The plasmid encoding the mutant T7 RNA polymerase_Δ172-173_, which does not incorporate non-templated nucleotides to the 3’-end of RNA, was a kind gift from Dr John Perona of Portland State University, and the His-tagged enzyme was purified as previously described (43).

### Cloning and expression of the gene encoding *Sa*TrmK

The DNA encoding *Sa*TrmK with a TEVP-cleavable N-terminal His-tag, purchased as a g-block (IDT Integrated DNA Technologies), was amplified by the polymerase chain reaction (PCR) using the forward primer 5’-GGAATTCCATATGCACCATCATCATCACCAC-3’ and the reverse primer 5’-CCCAAGCTTTCACAGCACACGCTCAATTA-3’. The g-block was codon-optimised for expression in *Escherichia coli*. The amplified DNA fragment was digested with *Nde*I and *Hin*dIII restriction enzymes and ligated into *Nde*I/*Hin*dIII-linearized pJexpress411 vector. The resulting plasmid was sequenced (Eurofins Genomics) to confirm insertion of the gene and that no mutation was introduced. The pJexpress411-*Sa*TrmK construct was transformed into *E. coli* BL21(DE3) competent cells. The transformed cells were grown in 1 L of lysogeny broth (LB) containing 50 μg mL^−1^ kanamycin at 37 °C to an optical density at 600 nm of 0.6. The culture was then equilibrated to 16 °C, and expression was induced with 1 mM IPTG. Cells were allowed to grow for an additional 20 h, harvested by centrifugation at 6,774 *g* for 15 min and stored at −20 °C.

### Purification of *Sa*TrmK

All purification procedures were carried out at 4 °C and chromatographic steps employed an AKTA Start FPLC system (GE Lifesciences). Cells were allowed to thaw on ice for 20 min before being re-suspended in buffer A (50 mM HEPES, 10 mM imidazole, 300 mM NaCl, pH 8.0) containing 0.2 mg mL^−1^ lysozyme, 0.05 mg mL^−1^ DNAse I, and half a tablet of EDTA-free Complete protease inhibitor cocktail, disrupted in a high-pressure cell disruptor (Constant Systems), and centrifuged at 48,000 *g* for 30 min to remove cell debris. The supernatant was filtered through a 0.45-μm membrane and loaded onto a HisTrap FF 5 mL column (GE Healthcare) pre-equilibrated with buffer A. The column was washed with 10 column volumes (CV) of buffer A, and the adsorbed proteins were eluted with 20 CV of a linear gradient of 0% to 80% buffer B (50 mM HEPES, 300 mM imidazole, 300 mM NaCl, pH 8.0). Fractions containing the desired protein were pooled and dialysed twice against 2 L of buffer C (20 mM HEPES, 150 mM NaCl, 2 mM DTT, 10% glycerol (*v/v*), pH 7.5). After dialysis, the protein was mixed with TEVP at a ratio of 1.3 mg of TEVP to 10 mg of *Sa*TrmK. The protein was further dialysed twice against 2 L of buffer C, then once against 2 L of buffer A. Samples were filtered through a 0.45-μm membrane and loaded onto a HisTrap FF 5 mL column (GE Healthcare) pre-equilibrated with buffer A. The flow through was collected and analysed by SDS-PAGE (NuPAGE Bis-Tris 4-12% Pre-cast gels, ThermoFisher Scientific), concentrated using 10,000 molecular weight cut off (MWCO) ultrafiltration membranes (Millipore), dialysed twice against 2 L of 20 mM HEPES pH 8.0, aliquoted, and stored at −80 °C. The concentration of *Sa*TrmK was determined spectrophotometrically (NanoDrop) at 280 nm using the theoretical extinction coefficient (ε_280_) of 15,930 M^−1^ cm^−1^ (Expasy) (44). The molecular mass of the protein, which contains an N-terminal glycine residue left following TEVP cleavage, was determined by electrospray-ionisation/time-of-flight (ESI/TOF) mass spectrometry (MS).

### *Sa*TrmK oligomeric state determination

Size-exclusion chromatography was carried out at 20 °C on a Superdex 200 10/300 GL column pre-equilibrated with 20 mM HEPES pH 7.5, using an AKTA Purifier FPLC system (GE Healthcare). Samples (1 mL) were loaded onto the column at 1 mg/mL. Vitamin B12 (1,350 Da), horse myoglobin (17,000 Da), chicken ovalbumin (44,000 Da), bovine γ-globulin (15,8000 Da) and bovine thyroglobulin (670,000 Da) (Bio-Rad) were used as MW standards. The logarithm of the MW of the standards were plotted against the ratios of the respective elution volumes (v_e_) to the void volume (v_0_). The points were then fitted to a linear regression and the values of slope and intercept were used to determine the molecular weight of *Sa*TrmK in solution.

### Crystallisation of *Sa*TrmK

All crystals were grown at 20 °C using *Sa*TrmK (10 mg mL^−1^) in 20 mM HEPES pH 8.0. Sitting drops were set up by mixing 150 nL of protein with 150 nL of in-house and commercial stochastic screens. Diffracting crystals were obtained with 0.1 M ammonium acetate, 0.1 M Bis-Tris pH 5.5, 17% PEG 10,000. For SAM and SAH co-crystallisation, the same condition was used, but *Sa*TrmK was incubated with either SAM or SAH (1 mM) for 1 h prior to sitting drops being set up.

### X-ray diffraction data collection and processing

X-ray diffraction data were collected on beamline I04 at Diamond Light Source, UK, and processed and scaled using the automated processing pipeline with Xia2 (45) using either DIALS (46) or XDS (47). The structures were solved by molecular replacement using PhaserMR (48) using Protein Data Bank (PDB) entry 3KR9 (7) as the search model for the *Sa*TrmK apoenzyme structure, which in turn was used as the search model for the SAH- and SAM-bound structures. Structures were refined using cycles of model building with COOT (49) and refinement with Refmac (50) and Phenix (51). Where clear electron density was observed, ligands (SAM, SAH, as well as glycerol, and citrate for the apoenzyme) were inserted in COOT using the existing coordinates and dictionaries. The structures were validated using MolProbity (52).

### MD simulations

MD simulations were carried out for *Sa*TrmK apoenzyme and the SAM- and SAH-bound binary complexes using the AMBER99SB force field as implemented in GROMACS (53) using the respective crystal structures as starting points. To prepare the system for simulations, each structure was placed in a cubic box at 8 Å from the box boundary, and solvated with TIP3P water model (54) then neutralized with potassium ions. Principal component analysis was carried out with gmx covar and gmx anaeig as implemented in GROMACS. Motional correlation studies used Wordom (55). Electrostatics were calculated using the Particle-Mesh Ewald sums method (56) with a real space cut-off of 10 Å, using order of 4 and a relative tolerance between long- and short-range energies of 10^−5^. Short range interactions were evaluated using a neighbour list of 10 Å and the Lennard-Jones interactions and the real space electrostatic interactions were truncated at 9 Å. The temperature was maintained at 300 K using V-rescale; hydrogen bonds were constrained using the LINCS algorithm. Energy minimization was carried out to reach a maximum force of no more than 10 kJ/mol using steepest descent algorithm. Mulliken charges of SAM and SAH were calculated for structures optimised by density-functional theory at B3LYP/6-31G(d) as implemented in GAMESS (57). The topology of SAM and SAH was defined with ACPYPE. Data reported are the mean of two independent simulations. Each structure was simulated twice for 500 ns, totalling 3 μs of simulation time. Electrostatic potential surfaces were calculated with the PDB2PQR pipeline (58) and snapshots taken every 125 ns of simulation.

### *In-silico* screening of *Sa*TrmK-binding compounds

Simple chemical scaffolds such as benzene, pyridine, purine and indole were used to explore the thermodynamic fit to *Sa*TrmK:SAM complex via *in-silico* molecular docking to ensemble conformations of *Sa*TrmK:SAM generated by normal mode analysis. Docking studies were conducted using an in-house proprietary 7D-Grid technology that applies a geometric search algorithm to generate ligand conformations in the protein structure, which are then placed in a grid composed of quantum polarised probes to calculate the binding energies. Energy maps were computed using the fragment molecular orbital method (59). The scaffold that produced the best docked conformation was used as a template in a 3D-structure search against an in-house phytochemical database. The phytochemical database was composed of derivatives of azadirachtin from the neem tree, curcumin from the turmeric plant, cucurbitacin from squash, chlorogenic acids and naphthoquinones from various plants.

### *In-vitro* transcription

DNA encoding for *S. aureus* tRNA^Leu^, A22C-tRNA^Leu^, A22G-tRNA^Leu^, A22U-tRNA^Leu^ or for small (18mer) RNA hairpins (RNA^18mer^, A10G-RNA^18mer^, A11U-RNA^18mer^, U5C/A14G-RNA^18mer^, A11-RNA^18mer^, and 0A-RNA^18mer^), all containing a T7 promoter sequence at the 5’-end, were amplified using primers (IDT) listed in Table S2. PCR products were used as templates for *in-vitro* transcription reactions. All *in-vitro* transcriptions were carried out under RNAse-free conditions. Solutions were prepared in diethyl pyrocarbonate (DEPC)-treated water. To set up a 1-mL transcription reaction, the following components were mixed in *in-vitro* transcription buffer (40 mM Tris-HCl pH 8.0, 22 mM MgCl_2_, 0.2 mM spermidine, 20 mM DTT and 0.1 μL Triton X-100) prior to incubation at 37 °C for 4 h: 20 μg DNA template, 5 mM ATP, 5 mM CTP, 5 mM UTP, 6 mM GTP, 7.8 μM T7 RNA polymerase. The reaction was spun down to remove pyrophosphate. RQ1 RNAse-free DNAse (Promega) was added to the supernatant, followed by incubation at 37 °C for at least 2 h. A phenol/chloroform extraction was carried out followed by an ethanol precipitation using 80 μL of 500 mM EDTA, 100 μL of 250 mM NaCl and 9 mL of 100% ethanol. The RNA was left to precipitate at −80 °C for at least 1 h and then centrifuged at 6,774 g for 1 h. The pellet was washed with 70% ethanol, dried and resuspended in storage buffer (10 mM MOPS pH 6.0 and 1 mM EDTA). RNAs were visualised in Novex TBE-Urea 15% Pre-cast gels (ThermoFisher Scientific). Prior to use in any experiment, RNAs were desalted in a Bio-Rad Micro Bio-Spin column, dialysed in Slide-A-Lyzer cassettes (Thermo Fisher Scientific) (2,000 MWCO for RNA^18mer^; 10,000 MWCO for tRNA) against 200 mL of DEPC-treated water, unfolded by heating at 95 °C for 10 min, and refolded by slowly cooling down at room temperature for 1 h. RNA concentration was determined spectrophotometrically at 260 nm. Secondary structure predictions for RNA^18mer^ were performed with RNAstructure (60).

### *Sa*TrmK activity assay and steady-state kinetics

A Tecan Infinite Lumi plate reader was utilised for all *Sa*TrmK activity measurements. The MTase-Glo™ Methyltransferase Assay Kit (Promega) was used to monitor the methyltransferase activity of *Sa*TrmK. This discontinuous assay converts the reaction product SAH to ATP, which is used in a luciferase reaction, and luminescence is detected. Luminescence is correlated to SAH concentration using an SAH standard curve (0 – 2 μM SAH) determined under the same conditions as the activity assay; 16 nM was the lowest SAH concentration reliably detected under these conditions. All assays were performed at 20 °C. Unless otherwise stated, a typical reaction mixture (20 μL) contained 100 mM HEPES pH 7.5, 3 mM MgCl_2_, 50 nM *Sa*TrmK, 0.5 μM tRNA or RNA^18mer^, and 0.5 μM SAM. For the dependence of SAH formation on *Sa*TrmK concentration, 0 – 150 nM enzyme was used. Inhibition by sinefungin was determined in the presence of 250 nM *Sa*TrmK, 2 μM tRNA, 2 μM SAM in presence or absence of 10 μM sinefungin. For the effect of sinefungin on the assay, 2 μM SAH replaced SAM, and *Sa*TrmK was omitted. Inhibition by plumbagin was determined in the presence of 250 nM *Sa*TrmK, 2 μM tRNA, 2 μM SAM, 1% DMSO in presence or absence of 5 μM plumbagin. For the effect of plumbagin on the assay, 2 μM SAH replaced SAM, and *Sa*TrmK was omitted. For reactions with A11-RNA^18mer^ and 0A-RNA^18mer^ as substrates, 1 μM *Sa*TrmK, 20 μM RNA, and 20 μM SAM were used. All reactions were incubated for 20 min, after which 5 μL MTase-Glo reagent was added and the reaction was incubated for 30 min. Finally, 25 μL of the MTase-Glo detection solution was added and the reaction incubated for an additional 30 min prior to luminescence counting. Control experiments lacked *Sa*TrmK, and their signal was subtracted from the corresponding reaction signal. In the instances where the effect of inhibitor on the assay itself was tested, where *Sa*TrmK was omitted, controls lacked SAH, and their signal was subtracted from the corresponding reaction signal with SAH. All measurements were performed at least in duplicate.

### Detection of N^1^-methyl-AMP by LC-MS

Reactions (50 μL) for methylation of tRNA were carried out in 100 mM HEPES pH 7.5, 3 mM MgCl_2_, with 20 μM *Sa*TrmK, 50 μM SAM and 50 μM either tRNA^Leu^ or A22C-tRNA^Leu^, and incubated at 20 °C for 2.5 h. The reaction (50 μL) for methylation of A11-RNA^18mer^ was carried out in 100 mM HEPES pH 7.5, 3 mM MgCl_2_, with 50 μM *Sa*TrmK, 150 μM SAM and 150 μM A11-RNA^18mer^, incubated at 20 °C for 1 h, after which another 50 μM *Sa*TrmK was added and the reaction was incubated at 20 °C for 1.5 h. For all experiments, controls lacking *Sa*TrmK were performed under the same conditions. To each sample, 0.1 volumes of 3 M sodium acetate and 3 volumes of ethanol were added. Samples were incubated overnight at −80 °C and subsequently centrifuged at 11,400 *g* for 10 min. The pellet was dissolved in 44 μL of water, to which 5 μL of 500 mM sodium acetate pH 5.5 and 1 μL of nuclease P1 (100 U) were added, and the reaction was incubated at room temperature for 10 min, followed by addition of [^15^N5]AMP at 10 μM as an internal standard, and methanol at a final concentration of 80%. Samples were incubated at −80 °C for 15 min and centrifuged at 11,400 *g* for 10 min. The supernatant was collected, dried under N2 and reconstituted with 50 μL of water. Commercial N^1^-methyl-AMP (10 μM) was treated in the same manner starting with the nuclease P1 reaction, and used as a standard. LC-MS was performed using a Waters ACQUITY UPLC system coupled to a Xevo G2-XS QToF mass spectrometer equipped with an ESI source. The autosampler was maintained at 4 °C throughout. Samples (10 μL) were loaded onto an Atlantis Premier BEH C18 AX column (2.1 × 100 mm, 1.7 μm) (Waters) at 40 °C and separated in (A) 0.1% formic acid in water and (B) 0.1% formic acid in acetonitrile as mobile phase in the following sequence: 0 – 1 min 99% A and 1% B, 1 – 9 min linear gradient from 99% A and 1% B to 1% A and 99% B, 9 – 11 min 1% A and 99% B, 11 – 15 min 99% A and 1% B at a flow rate of 400 μL min^−1^. Ion counts of the eluents were detected with a capillary voltage of 2.5 kV in positive ion mode. The source and desolvation gas temperatures of the mass spectrometer were set at 120 °C and 500 °C, respectively. The cone gas flow was set to 50 L/h, while the desolvation gas flow was set at 800 L/h. An MS^E^ scan was performed between 50 – 700 m/z. A lockspray signal was measured and a mass correction was applied by collecting every 10s, averaging 3 scans of 1 s each, using Leucine Enkephalin as a correction factor for mass accuracy.

### *Sa*TrmK thermal denaturation by DSF

DSF measurements (λ_ex_ = 490 nm; λ_em_ = 610 nm) were performed in 96-well plates on a Stratagene Mx3005p instrument. Thermal denaturation assays (50 μL) of 7 μM *Sa*TrmK were carried out in 500 mM HEPES pH 7.5 in the absence and in the presence of either 200 μM SAM, 10 μM SAH, or 0 – 640 μM citrate. Sypro Orange (5×) (Invitrogen) was added to all wells. Thermal denaturation curves were recorded over a temperature range of 25-93 °C with 1 °C min^−1^ increments. Control curves without the enzyme were subtracted from curves containing the enzyme. All measurements were carried out in triplicate. Data were fitted to equation 1 (61), where *F*U is fraction unfolded, *T* is the temperature in °C, *T*_m_ is the melting temperature, *c* is the slope of the transition region, and *LL* and *UL* are folded and unfolded baselines, respectively.

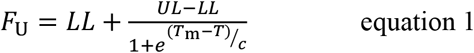

### Equilibrium binding by ITC

ITC measurements were carried out at 20 °C in a MicroCal PEAQ-ITC calorimeter (Malvern Instruments). *Sa*TrmK and ligands (either SAM or SAH) were solubilised in 100 mM HEPES pH 7.5. An initial injection of 0.4 μL was performed, followed by 18 successive injections of 2 μL of ligand (either 1 mM SAM or 209 μM SAH) into 300 μL of either 40 μM *Sa*TrmK (for SAM binding) or 35 μM *Sa*TrmK (for SAH binding) with 150-s interval between successive injections and a reference power of 10 μcal s^−1^. The heat of dilution for each experiment was measured by titrating SAM or SAH into 300 μL assay buffer and subtracted from the corresponding binding curve. All measurements were performed in duplicate. Binding curves were fitted to a single-site binding model as implemented in the PEAQ-ITC analysis software (Malvern Instruments). All values reported represent the mean ± standard error. One of the injections of SAM yielded an outlier, likely due to an air bubble, and was excluded from the fit.

### *Sa*TrmK intact mass and peptide analysis in the presence of plumbagin

*Sa*TrmK (20 μM) was incubated in 100 mM HEPES pH 7.5, 3 mM MgCl_2_, 1% DMSO in the presence and absence of 30 μM plumbagin at 20 °C for 20 min. Samples were diluted to obtain 1 μM protein in eluent A (95% water, 5% acetronitrile, 1% formic acid), from which 10 μL were loaded onto a Waters MassPREP desalting cartridge (2.1 mm × 10 mm) and analysed by LC-MS using a Waters ACQUITY UPLC coupled to a Xevo G2 TOF mass spectrometer with Masslynx software. A linear gradient from 95% A and 5% B (5% water, 95% acetronitrile, 1% formic acid) to 5% A and 95% B over 6 min, before returning to 95% A and 5% B, was performed at flow rate of 200 μL min^−1^. MS data were collected from 500 – 2,500 m/z. The charged ion series was deconvoluted to 0.1-mass-unit-resolution using MaxEnt1 deconvolution algorithm using a full width at half maximum of 0.4 m/z. For proteolytic digestion analysis, *Sa*TrmK was incubated with and without plumbagin under the same conditions described above. Samples were diluted to obtain 10 μM protein in 10 mM ammonium bicarbonate. To each 10-μL sample, 0.1 μg of trypsin was added and incubated at 30 °C for 8 hours, followed by addition of 0.1 μg of Glu-C and a further incubation for 8 hours. Reactions were acidified and analysed by LC-MS/MS on an Eksigent 2D ultra nanoLC coupled to a Sciex 5600+ mass spectrometer set up in trap-elute format. The sample was loaded onto a Thermoscientific 2-cm PepMap™ trap column and washed with 0.05% trifluoracetic acid for 5 min at 5 μL min^−1^; the trap was then switched in-line to the Thermoscientific PepMap^TM^ analytical column (75 μm × 15 cm), and a linear gradient of 0.1% formic acid to 0.1% formic acid and acetonitrile over 1 h applied. The eluting peptides were sprayed directly into the mass spectrometer. For discovery proteomics, the mass spectrometer was operated in data-dependent-mode scanning MS from 400 – 1200 m/z and the top-10 most intense peptides with charge states between 2+ and 5+ were selected for CID fragmentation with MS/MS data collected from 95 – 1800 m/z. The raw data were analysed in Mascot 2.6 (Matrix Science) using the msconvert data extraction script (62). Data were searched against an in-house database of proteins to which the *Sa*TrmK sequence was added using trypsin/Glu-C combined as digestion enzyme with error tolerant search settings to include any possible modification. For confirmation of the peptide modification, the mass spectrometer was set to carry out MS scan on 400 – 1200 m/z and dedicated product ion scan fragmentation on [M+2H]^2+^ = 923.4972 and [M+2H]^2+^ = 1,016.5033 collecting MS/MS data from 95 – 2000 m/z after CID at 45 V. Data were displayed according to Biemann’s notation (63).

## Data availability

Atomic coordinates and structure factors for the reported crystal structures have been deposited with the Protein Data bank under accession numbers 7O4M, 7O4N, and 7O4O. Molecular dynamics and docking atomic coordinates are held at Kcat Enzymatic Private Limited and may be available upon request to Dr Pravin Kumar. All other data presented are contained within the manuscript.

## Supporting information

This article contains supporting information.

## Acknowledgements

X-ray diffraction data were collected at Diamond Light Source, UK, on beamline I04. The authors thank Dr Magnus S. Alphey for his assistance with the initial crystallisation trial for *Sa*TrmK, and Dr John Perona for his kind gift of the mutant T7 RNA polymerase-encoding plasmid.

## Author contributions

PS, AC, CJL and GF collected all molecular biology, biochemistry, and biophysics data. VO and TMG carried out crystallographic data collection and analysis. AK, DR, NBK, RL, LM, GS and PK performed all MD simulations and *in-silico* screening. ES and CMC collected and analysed smallmolecule MS data. SS and SLS collected and analysed protein MS data. RGdS conceived and directed the research. RGdS, PS, TMG, ES and PK wrote the manuscript.

## Funding and additional information

This work was supported by a Wellcome Trust Seed Award in Science [208980/Z/17/Z] to RGdS; a University of St Andrews/Scottish Funding Council St Andrews Restarting Research Fund to RGdS; and a Wellcome Trust Institutional Strategic Support Fund [204821/Z/16/Z] to the University of St Andrews. ES is the recipient of a Cunningham Trust PhD studentship (PhD-CT-18-41).

## Conflict of interest

The authors declare that they have no conflicts of interest with the contents of this article.

## Abbreviations

m^1^A: N1-methyl-adenine
TrmK: m^1^A22-tRNA methyltransferase
*Sa*TrmK: *S. aureus* TrmK
ITC: isothermal titration calorimetry
MD: molecular dynamics
DSF: differential scanning fluorimetry
ESI: electrospray-ionisation
*T*_m_: melting temperature
H-bond: hydrogen bond
TEVP: tobacco etch virus protease

## Notes

### Competing Interest Statement

The authors have declared no competing interest.

