## Supporting information for "Structure, dynamics, and inhibition of *Staphylococcus aureus* m^1^A22-tRNA methyltransferase"

**List Of Supporting Figures and Tables:**

**Figure S1. SDS-PAGE analysis of *Sa*TrmK purification**
**Figure S2. ESI-TOF-MS analysis of purified *Sa*TrmK**
**Figure S3. DSF-based thermal denaturation assay of *Sa*TrmK**
**Figure S4. Determination of the oligomeric state of *Sa*TrmK in solution by analytical gel filtration**
**Table S1. X-ray data processing and refinement statistics**
**Figure S5. Overlay of *Sa*TrmK, *Sa*TrmK:SAM, and *Sa*TrmK:SAH over all Cα atoms**
**Figure S6. Overlay of Cα atoms of *Sa*TrmK, *B. subtilis* TrmK (PDB entry 6Q56), and *S. pneumoniae* TrmK (PDB entry 3KR9)**
**Figure S6. Overlay of Cα atoms of *Sa*TrmK, *B. subtilis* TrmK (PDB entry 6Q56), and *S. pneumoniae* TrmK (PDB entry 3KR9)**
**Figure S7. Possible interaction between *Sa*TrmK and citrate**
**Figure S8. Position of Asp26 side chain in *Sa*TrmK:SAM and *Sa*TrmK apoenzyme**
**Figure S9. Domain cross-correlation matrices**
**Figure S10. Molecular electrostatic potential surfaces**
**Figure S11. LC-MS for SAH detection**
**Figure S12. Dependence of luminescence counts on SAH concentration**
**Figure S13. Effect of sinefungin on the quantification of SAH via the MTase-Glo™ Methyltransferase Assay**

**Figure S14. Effect of plumbagin on the quantification of SAH via the MTase-Glo™ Methyltransferase Assay**

**Figure S15. Potential covalent adducts of plumbagin with *Sa*TrmK**

**Table S2. Oligonucleotide primers for production of DNA templates for *in-vitro* transcription**

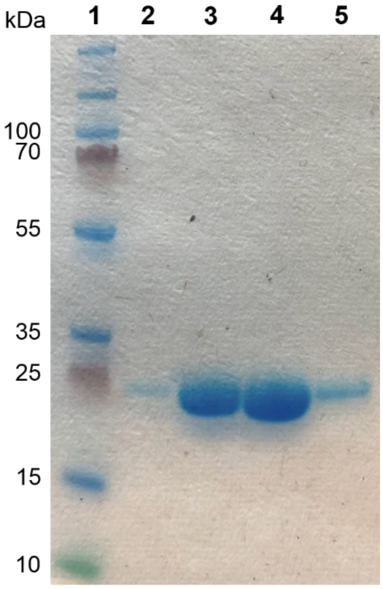

**Figure S1. SDS-PAGE analysis of *Sa*TrmK purification.** Lane 1, Thermo Scientific Page Ruler Plus Prestained Protein Ladder. Lane 2 – 5, flowthrough from the HisTrap FF 5 mL column after TEVP cleavage.

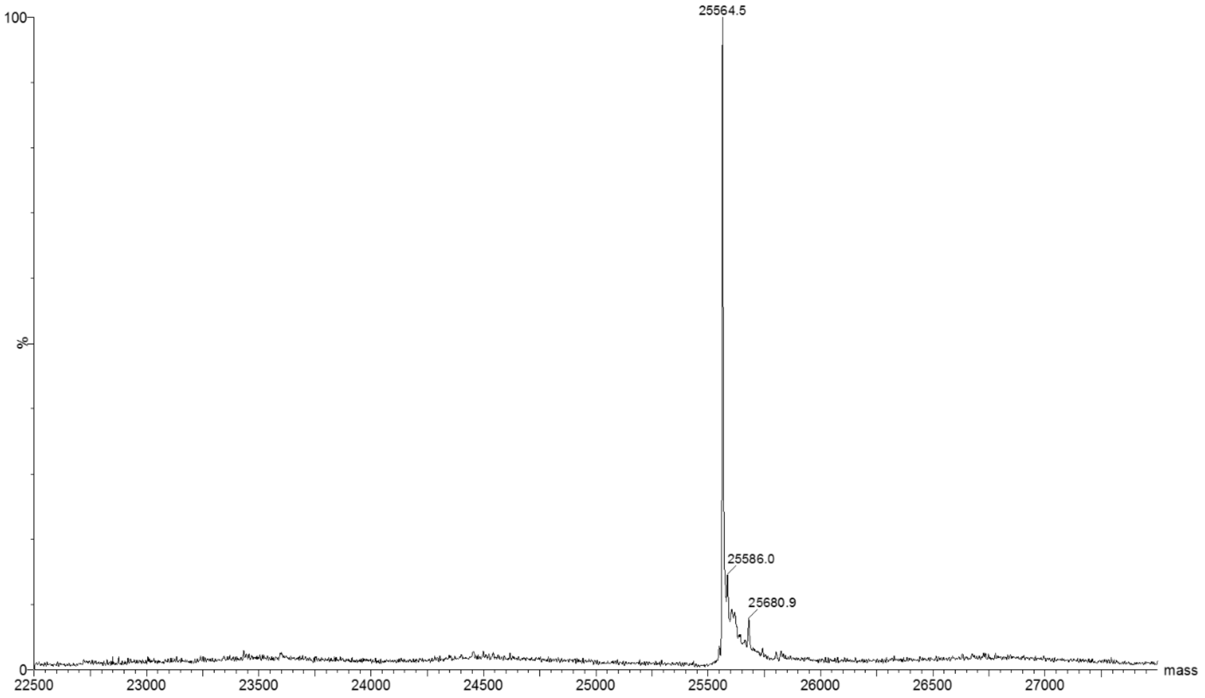


**Figure S2. ESI-TOF-MS analysis of purified *Sa*TrmK.** The experimental mass matches the predicted molecular mass of 25,565.3.

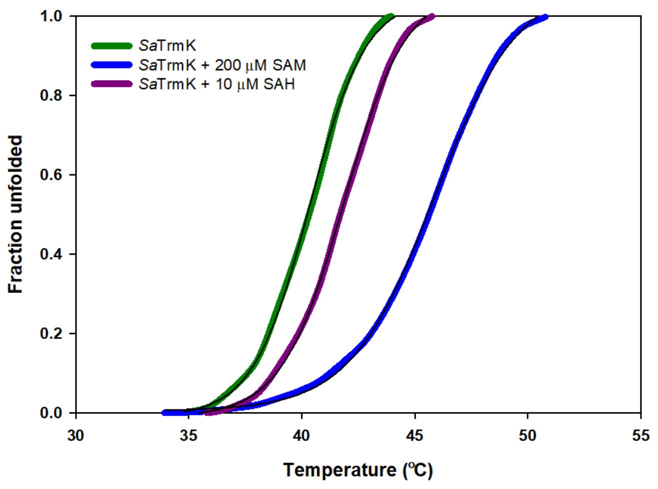

**Figure S3. DSF-based thermal denaturation assay of *Sa*TrmK.** Lines of best fit to equation 1 are in black.

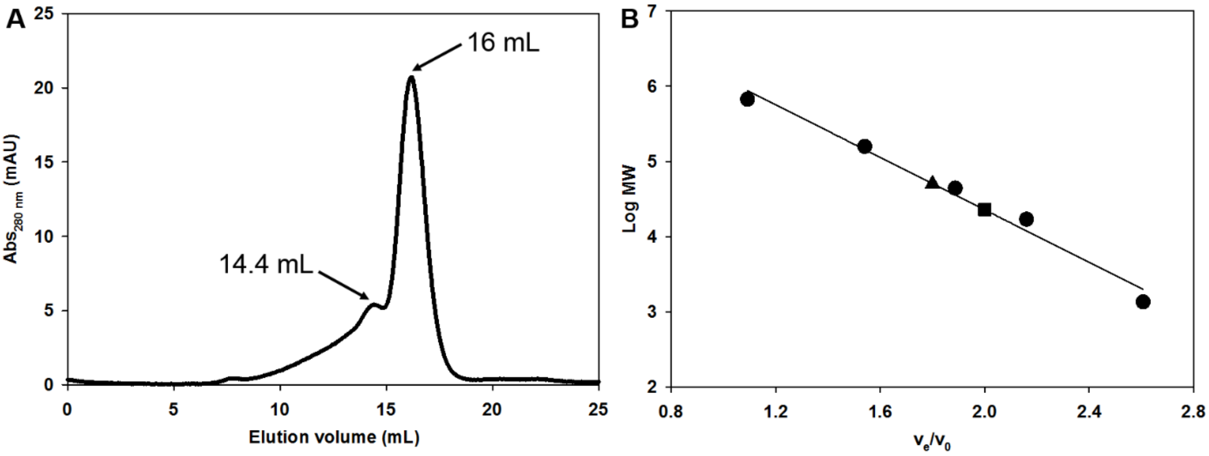


**Figure S4. Determination of the oligomeric state of *Sa*TrmK in solution by analytical gel filtration. (A)** Elution profile of *Sa*TrmK. **(B)** Relationship between molecular weight and elution volume ($v_e$) to void volume ($v_0$) ratio using molecular weight standards (circles). The line is a linear regression of the data. The $v_e/v_0$ ratio involving the major and minor elution volumes of *Sa*TrmK are plotted as square and triangle, respectively.

**Table S1. X-ray data processing and refinement statistics.**

| | **TrmK apo** | **TrmK:SAM** | **TrmK:SAH** |
|---|---|---|---|
| **PDB** | **7O4M** | **7O4N** | **7O4O** |
| **Wavelength (Å)** | 0.9795 | 0.9795 | 0.9795 |
| **Resolution range (Å)** | 43.8 - 1.09 (1.13 - 1.09) | 43.4 - 1.40 (1.45 - 1.40) | 19.4 - 1.52 (1.57 - 1.52) |
| **Space group** | $P\ 2_1\ 2_1\ 2_1$ | $P\ 2_1\ 2_1\ 2_1$ | $P\ 2_1\ 2_1\ 2_1$ |
| **Unit cell dimensions** | | | |
| **a, b, c (Å)** | 59.29, 60.72, 63.14 | 59.64, 61.31, 63.28 | 59.75, 61.44, 63.19 |

| | | | |
|---|---|---|---|
| **α, β, γ (°)** | 90, 90, 90 | 90, 90, 90 | 90, 90, 90 |
| **Total reflections** | 158738 (4396) | 91928 (8937) | 71273 (7033) |
| **Unique reflections** | 85297 (3451) | 46295 (4497) | 36225 (3577) |
| **Multiplicity** | 1.9 (1.2) | 2.0 (2.0) | 2.0 (2.0) |
| **Completeness (%)** | 88.7 (36.5) | 99.5 (98.8) | 99.3 (99.3) |
| **Mean I/sigma(I)** | 12.0 (0.93) | 13.38 (1.52) | 15.60 (1.54) |
| **R-merge** | 0.030 (0.61) | 0.039 (0.68) | 0.034 (0.48) |
| **CC1/2** | 0.99 (0.52) | 0.99 (0.44) | 0.99 (0.39) |
| **Reflections used in refinement** | 84861 (3442) | 46146 (4497) | 36225 (3576) |
| **R-work** | 0.139 | 0.148 | 0.169 |
| **R-free** | 0.170 | 0.199 | 0.231 |
| **RMSD (bonds)** | 0.012 | 0.011 | 0.0011 |
| **RMSD (angles)** | 1.66 | 1.61 | 1.70 |
| **Number of non-hydrogen atoms** | 2315 | 2197 | 2150 |
| **Protein** | 1921 | 1880 | 1839 |
| **Ligands** | 19 | 33 | 38 |
| **Solvent** | 375 | 284 | 273 |
| **Average B-factor ($Å^2$)** | 15.4 | 19.9 | 20.6 |
| **Protein** | 12.7 | 17.6 | 18.5 |
| **Ligands** | 16.6 | 22.8 | 19.9 |
| **Solvent** | 28.9 | 35.2 | 34.8 |

Values within brackets are for the highest resolution shell.

RMSD, root mean square deviation

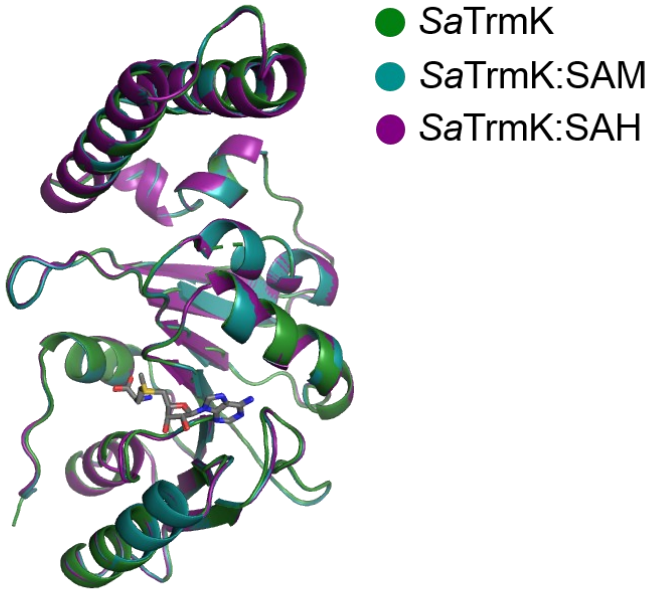


**Figure S5. Overlay of *Sa*TrmK, *Sa*TrmK:SAM, and *Sa*TrmK:SAH over all Cα atoms.**

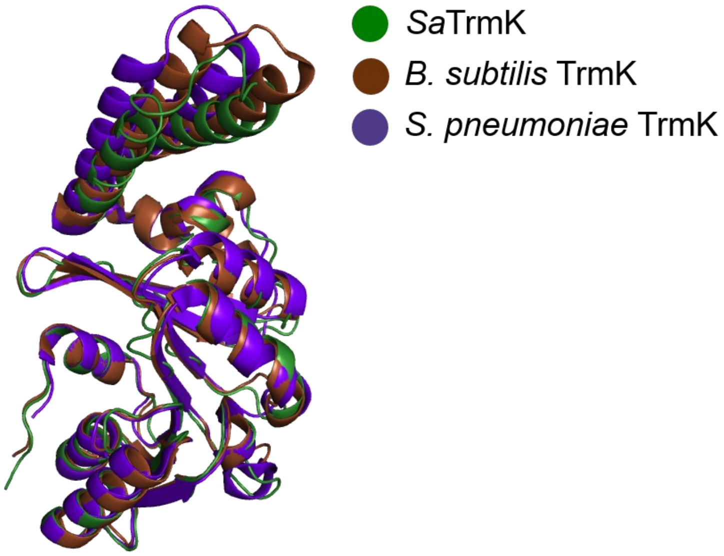


**Figure S6. Overlay of Cα atoms of *Sa*TrmK, *B. subtilis* TrmK (PDB entry 6Q56), and *S. pneumoniae* TrmK (PDB entry 3KR9).**

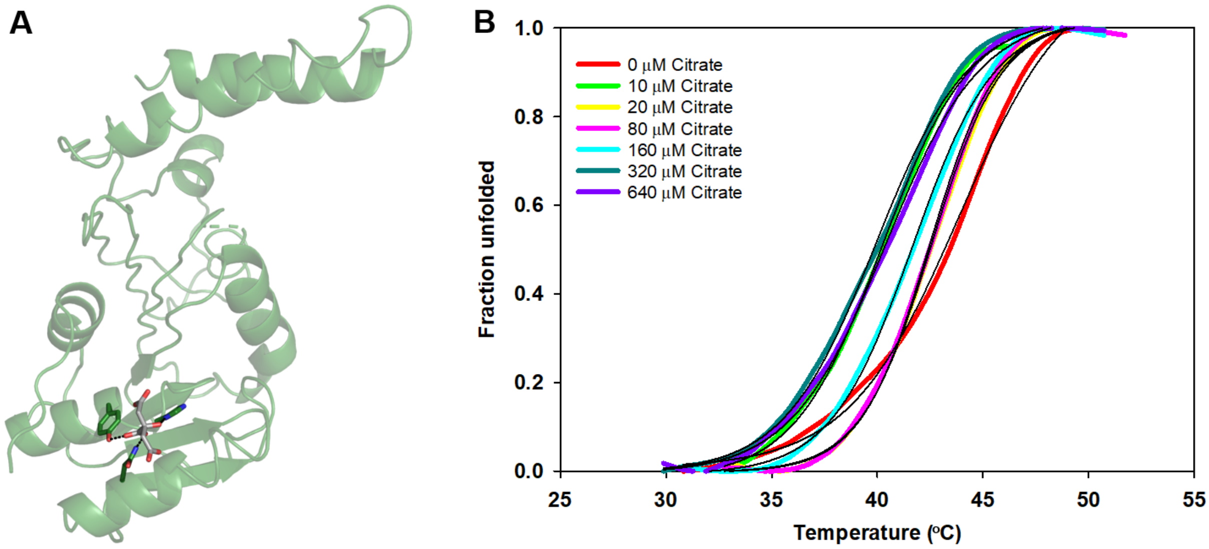


**Figure S7. Possible interaction between *Sa*TrmK and citrate. (A)** Polar contacts (dashed lines) among citrate and the side chains of His27, Tyr29, and Asn59. Side chains (green) and citrate (grey) are depicted as stick models. **(B)** DSF-based thermal denaturation assay of *Sa*TrmK in the presence and absence of citrate. Lines of best fit to equation 1 are in black, yielding $T_m$ (°C) of 43.0 ± 0.2, 40.2 ± 0.1, 42.5 ± 0.1, 42.04 ± 0.06, 41.67 ± 0.08, 39.9 ± 0.09, 40.4 ± 0.1, respectively, from 0 – 640 μM citrate.

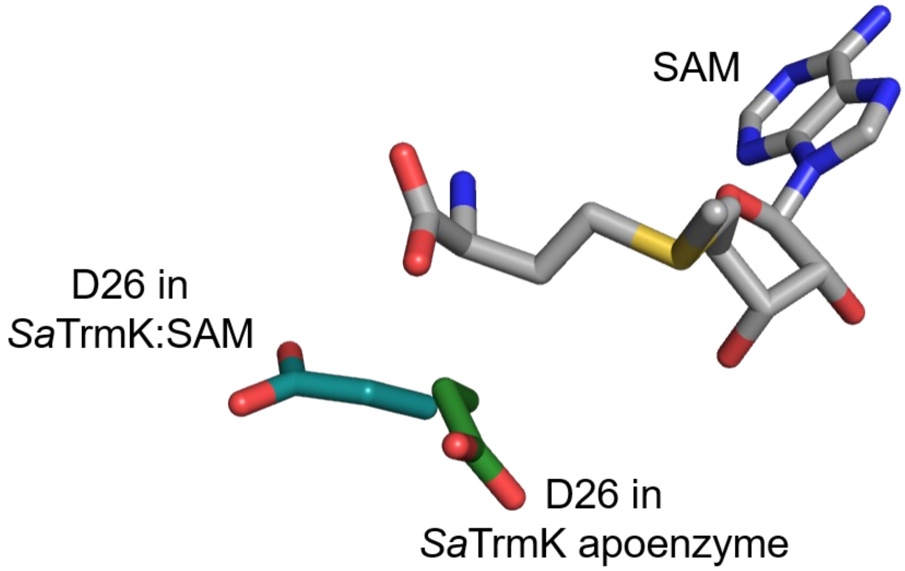


**Figure S8. Position of Asp26 side chain in *Sa*TrmK:SAM and *Sa*TrmK apoenzyme.** Asp26 side chain and SAM are shown in stick models.

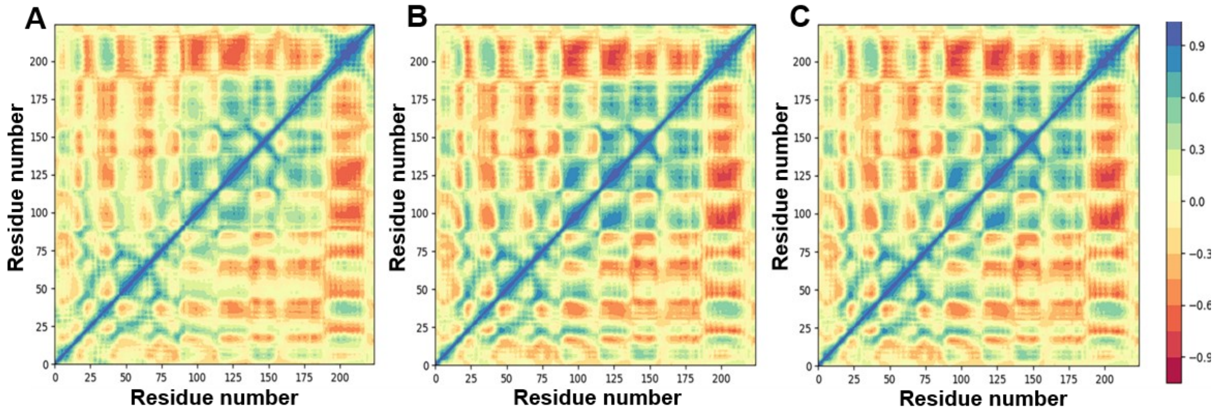


**Figure S9. Domain cross-correlation matrices. (A)** *Sa*TrmK apoenzyme. **(B)** *Sa*TrmK:SAM. **(C)** *Sa*TrmK:SAH.

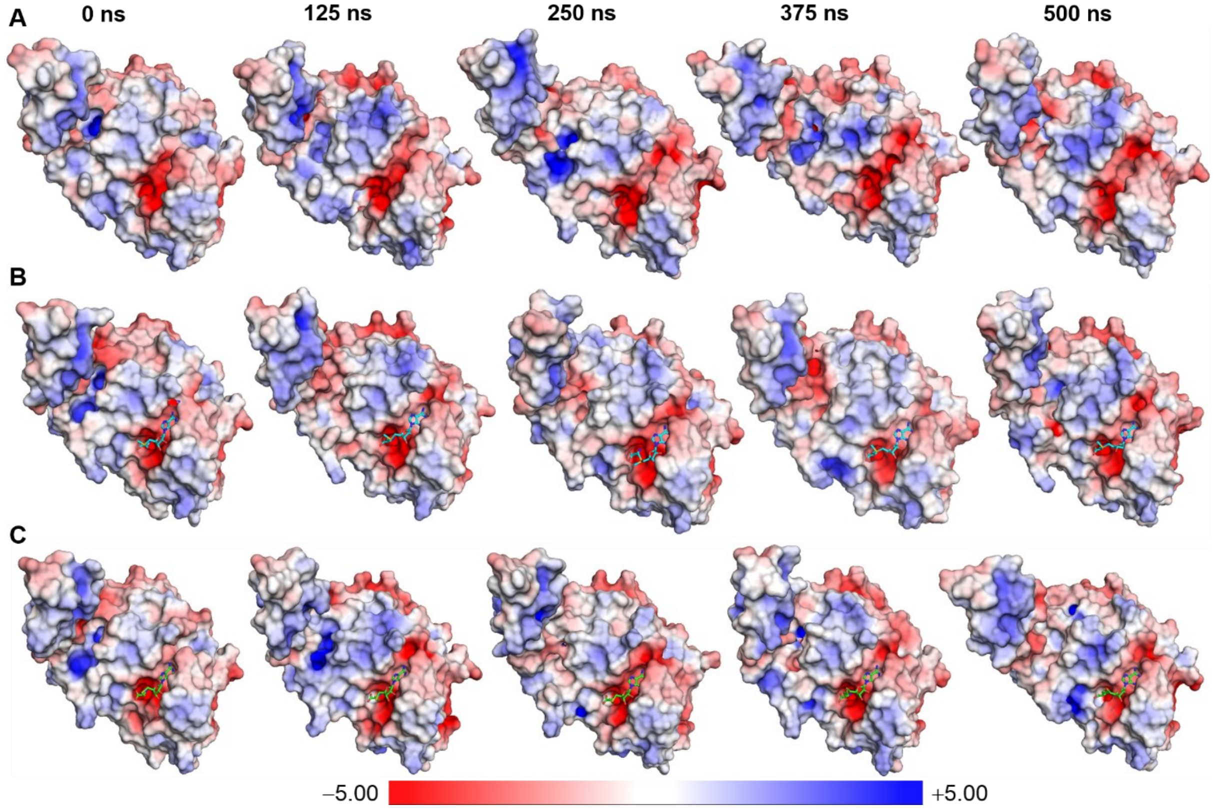


**Figure S10. Molecular electrostatic potential surfaces. (A)** Time-dependent molecular electrostatic potential surface of *Sa*TrmK. **(B)** Time-dependent molecular electrostatic potential surface of *Sa*TrmK:SAM, where SAM is shown as stick model. **(C)** Time-dependent molecular electrostatic potential surface of *Sa*TrmK:SAH, where SAH is shown as stick model. For all surfaces, snapshots of the electrostatic potential were taken at the times indicated at the top along the 500 ns of MD simulations. The units of the colour saturation are kT/e.

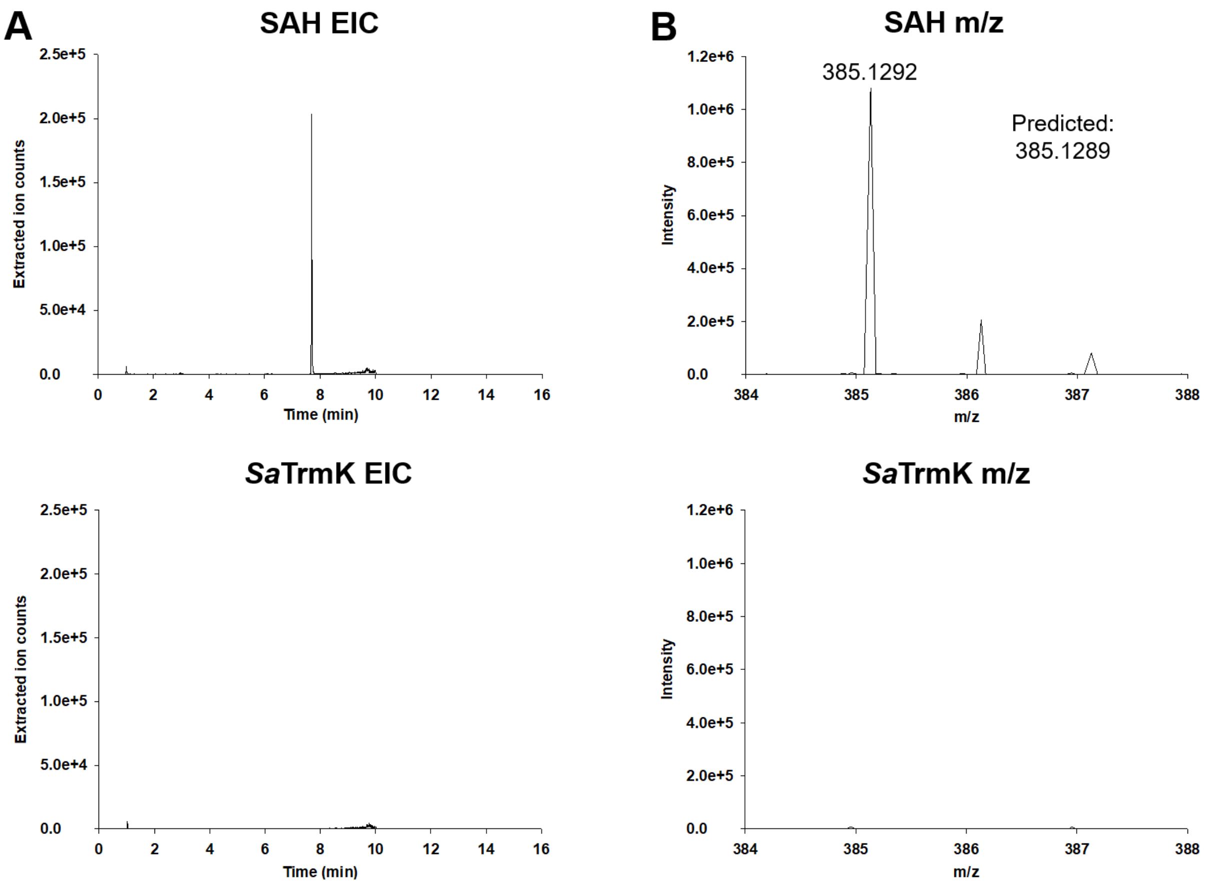


**Figure S11. LC-MS for SAH detection. (A)** Extracted ion counts of TCA-treated SAH and TCA-treated SaTrmK. **(B)** Detected mass from the respective extracted ion counts.

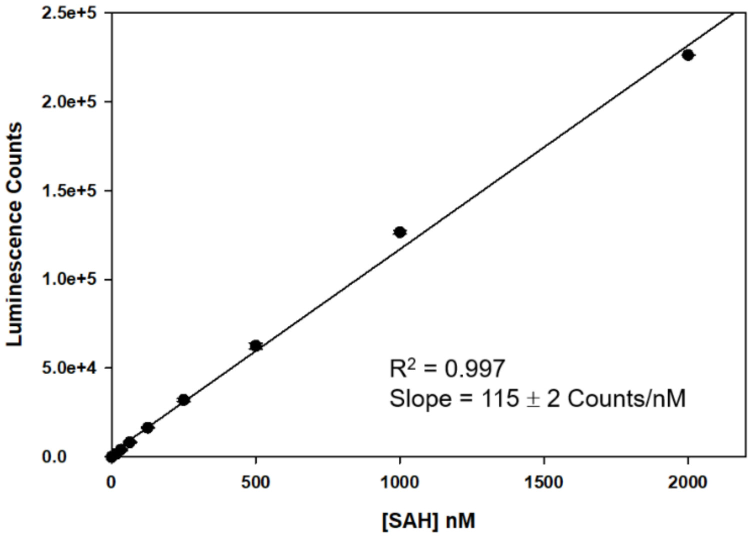


**Figure S12. Dependence of luminescence counts on SAH concentration.** Data are mean ± SD of duplicate measurements. The line is a linear regression of the data.

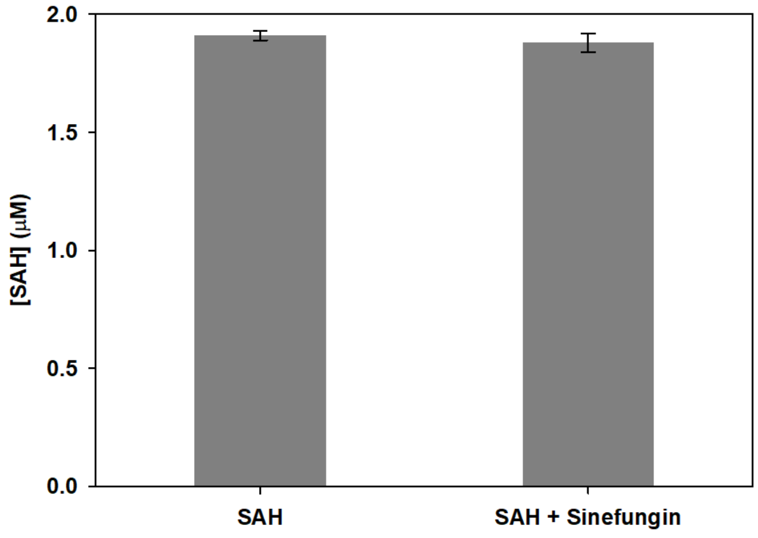


**Figure S13. Effect of sinefungin on the quantification of SAH via the MTase-Glo™ Methyltransferase Assay.** Data are mean ± SD of duplicate measurements.

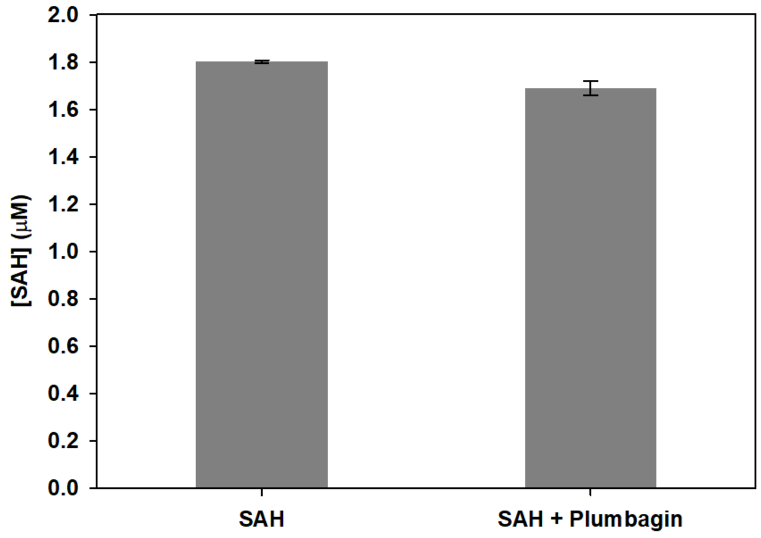


**Figure S14. Effect of plumbagin on the quantification of SAH via the MTase-Glo™ Methyltransferase Assay.** Data are mean ± SD of duplicate measurements.

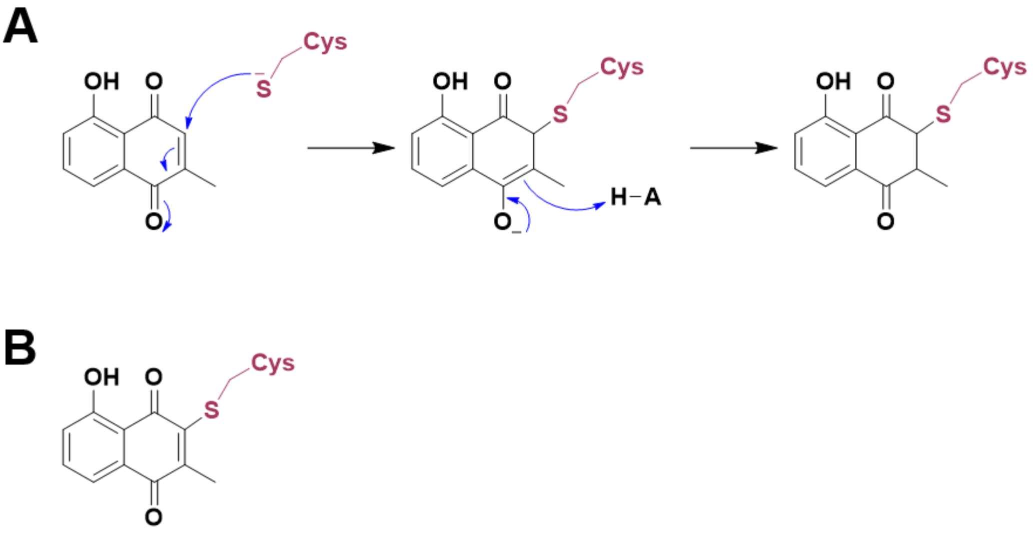


**Figure S15. Potential covalent adducts of plumbagin with *Sa*TrmK. (A)** The mechanism of Michael addition. The final Michael adduct would produce a mass difference of 188 in comparison with the unlabelled enzyme. **(B)** Putative oxidation product following Michael addition, which would result in a mass difference of 186 in comparison with the unlabelled enzyme.

**Table S2. Oligonucleotide primers for production of DNA templates for *in-vitro* transcription.**

| Amplicon | Primers (5′ to 3′) |
|---|---|
| **tRNA$^{Leu}$** | Forward Primer 1: CTCGAGTAATACGACTCACTATAGGCGGTCGTGGCGGAA<br>Reverse Primer 2: TGGTGGATGCGGCCGAG |
| **A22C-tRNA$^{Leu}$** | Forward Primer 1: CTCGAGTAATACGACTCACTATAGGCGGT |
| | Reverse Primer 2: AGGCCCTCAACCTAGCGCAGCTGCCATTCCGCCACGACCGCCTATAGTGAGTCGTATTA |
| | Forward Primer 3: GCTAGGTTGAGGGCCTAGTGGGAGAAGTCCCGTGGAGGTTCAAGTCCTCTCGGCCGCATC |
| | Reverse Primer 4: TGGTGGATGCGGCCGAGAG |
| **A22U-tRNA$^{Leu}$** | Forward Primer 1: CTCGAGTAATACGACTCACTATAGGCGGTCGTGGCGGAATGGCAGTTGCGCT |
| | Reverse Primer 2: CCCACTAGGCCCTCAACCTAGCGCAACTGCCATTC |
| | Forward Primer 3: TTGAGGGCCTAGTGGGAGAAGTCCCGTGGAGGTTCAAGTCCTCTCGGCCGCATC |
| | Reverse Primer 4: TGGTGGATGCGGCCGAGAG |
| **A22G-tRNA$^{Leu}$** | Forward Primer 1: CTCGAGTAATACGACTCACTATAGGCGGTCGTGGCGGAATGGCAGGTGCGCT |
| | Reverse Primer 2: CCCACTAGGCCCTCAACCTAGCGCACCTGCCATT |
| | Forward Primer 3: TTGAGGGCCTAGTGGGAGAAGTCCCGTGGAGGTTCAAGTCCTCTCGGCCGCATC |
| | Reverse Primer 4: TGGTGGATGCGGCCGAGAG |
| **RNA$^{18mer}$** | CTCGAGTAATACGACTCACTATAGGCGGTCGGCAACGACCGC |
| **A10G-RNA$^{18mer}$** | CTCGAGTAATACGACTCACTATAGGCGGTCGGCGACGACCGC |
| **A11G-RNA$^{18mer}$** | CTCGAGTAATACGACTCACTATAGGCGGTCGGCATCGACCGC |

| | |
|---|---|
| **U5C/A14G-RNA$^{18mer}$** | CTCGAGTAATACGACTCACTATAGGCGGCCGGCAACGGCCGC |
| **A11-RNA$^{18mer}$** | CTCGAGTAATACGACTCACTAGGCGGGCGGCGACGCCCGC |
| **0A-RNA$^{18mer}$** | CTCGAGTAATACGACTCACTAGGCGGGCGGCGGCGCCCGC |
